# Loss of the chromatin remodeler, *ATRX*, promotes aggressive features of osteosarcoma with increased NF-κB signaling and integrin receptor binding

**DOI:** 10.1101/2022.04.22.489086

**Authors:** Suzanne Bartholf DeWitt, Sarah Hoskinson Plumlee, Hailey E. Brighton, Dharshan Sivaraj, E. J. Martz, Maryam Zand, Vardhman Kumar, Maya U. Sheth, Warren Floyd, Jacob Spruance, Nathan Hawkey, Shyni Varghese, Jianhua Ruan, David G. Kirsch, Jason A. Somarelli, Ben Alman, William C. Eward

**Author notes:** **Declarations of Interests:** DGK is a co-founder of Xrad Therapeutics, which is developing radiosensitizers, and serves on the Scientific Advisory Board of Lumicell, which is commercializing intraoperative imaging technology. DGK also receives funding for a clinical trial from a Stand Up To Cancer (SU2C) Catalyst Research Grant with support from Merck. The laboratory of DGK currently receives funding or reagents from Xrad Therapeutics, Merck, Amgen, Bristol-Myers Squibb, Varian Medical Systems, and Calithera, but these did not support the research described in this manuscript.

## Abstract

Osteosarcoma (OS) is a lethal disease with few known targeted therapies. Here we show that decreased *ATRX* expression is associated with more aggressive tumor cell phenotypes, including increased growth, migration, invasion, and metastasis. These phenotypic changes correspond with activation of NF-κB signaling, extracellular matrix remodeling, increased integrin αvβ3 expression, and ETS family transcription factor binding. Here we characterize these changes *in vitro*, *in vivo*, and in a dataset of human OS patients. This increased aggression substantially sensitizes *ATRX*-deficient OS cells to integrin signaling inhibition. Thus, *ATRX* plays an important tumor suppression role in OS, and loss of function of this gene may underlie new therapeutic vulnerabilities. The relationship between *ATRX* expression and integrin binding, NF-κB activation, and ETS family transcription factor binding has not been described in previous studies and may impact the pathophysiology of other diseases with *ATRX* loss, including other cancers and the ATR-X alpha thalassemia mental retardation syndrome.

## Introduction

Osteosarcoma (OS) is the most common primary bone cancer diagnosed in humans and is most common among children and adolescents. It is a highly lethal cancer with a propensity for lung metastasis. At present, at least one-third of people diagnosed with OS die from this disease, even if a diagnosis is made and aggressive treatment with surgical excision and combination chemotherapy (i.e. doxorubicin, methotrexate, and cisplatin) is started early in the course of the disease (1, 2). Even for patients who do respond to therapy and survive OS, a normal life expectancy is unlikely due to the toxicity of current treatment regimens. For patients who develop metastatic disease, the prognosis is particularly bleak: almost 70% of patients do not survive beyond five years (3). Despite advances in understanding the molecular and genetic features underlying OS, patient outcomes have not improved significantly in over thirty years. Progress in OS research has been hindered, in part, by the rarity of the disease, with fewer than 1,000 human cases diagnosed annually in the United States (2), as well as by the genomic complexity of OS, with its considerable inter- and intra-tumoral heterogeneity (4–7). For all of these reasons, there remains an urgent need for a better understanding of the molecular underpinnings of OS biology and the resulting altered cellular pathways that may be targetable to provide more specific, refined, and effective therapies for OS.

Recent genomics studies have revealed a high rate of structural variation among OS tumors, including somatic mutations and copy number alterations, as well as a variety of single nucleotide variations or recurrent point mutations (5, 6, 8). Although the *TP53* and *RB1* genes show the most common recurrent alterations in OS, there are also commonly-recurrent somatic alterations in other candidate driver genes, such as *ATRX* (5, 9). In fact, across a variety of cancers, several recent studies have identified frequent loss-of-function mutations in *ATRX*, including in gliomas, pancreatic neuroendocrine tumors, melanomas and soft tissue sarcomas (10–16) (Figure 1A). The Cancer Genome Atlas identified *ATRX* as the 14^th^ most frequently altered gene across all cancers surveyed in the study (17–19). In 288 OS tumors surveyed by the American Association for Cancer Research Project Genomics Evidence Neoplasia Information Exchange (GENIE) Consortium, *ATRX* was the second most frequently mutated gene, second only to *TP53* (18–20) (Figure 1B). Additionally, at the protein level, several recent studies examining both human and canine OS tumors have found that between 20 and 30 percent lack nuclear expression of ATRX (11, 21, 22). Importantly, the actual incidence of *ATRX* mutation may be underestimated by modern next-generation sequencing technologies because these methods do not detect many complex indels with short sequence reads. Indeed, Ye et al. (23) identified *ATRX* as one of several oncogenic driver genes with frequent somatic complex indels in tumors across several cancer types that were overlooked by studies using next-generation sequencing modalities. Despite the known frequency and consistency across both human and canine OS, the impact of this *ATRX* loss on OS biology is not fully understood.

**Figure 1:**
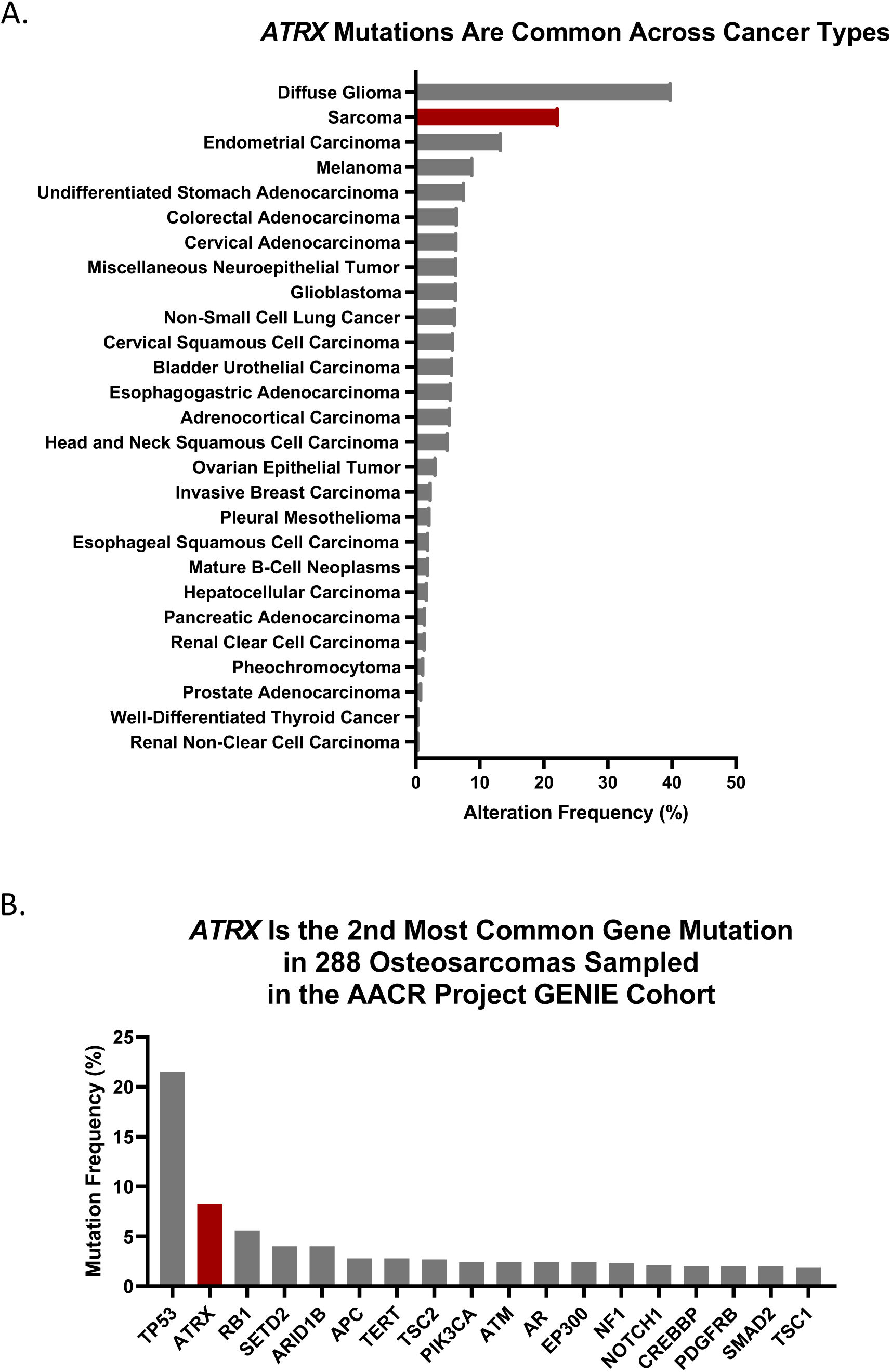
Frequency of *ATRX* mutations. A) Frequency of *ATRX* mutations across cancers in TCGA PanCancer Atlas, as accessed from cBioPortal (17–19). B) *ATRX* is the second most frequently mutated gene in 288 osteosarcomas surveyed by the AACR Project GENIE Consortium as accessed from cBioPortal (18–20).

*ATRX* (alpha thalassemia and mental retardation syndrome X-linked) is a member of the SWI/SNF family of chromatin remodeling factors. In humans, germline loss of function of the *ATRX* gene causes the <u>A</u>lpha <u>T</u>halassemia and mental <u>R</u>etardation, <u>X</u>-linked Syndrome, for which the gene is named (24). The ATRX protein contains two highly-conserved domains: the SWI/SNF helicase domain, which regulates chromatin remodeling, and the ATRX-DNMT3-DNMT3L (ADD) domain, which controls DNA methylation patterns and transcriptional repression (25, 26). With this ADD domain, ATRX forms dimers with death domain associated protein (DAXX), and this complex acts as a histone chaperone to deposit histone variant H3.3 to GC-rich regions of the genome, including the pericentric, ribosomal, and telomeric repeat sequences (27, 28). In the context of cancers, nearly all published studies have focused on the correlation between loss of *ATRX* expression and activation of the alternative lengthening of telomeres (ALT) pathway for telomere maintenance (21, 22, 29, 30). However, while replicative immortality is one of the key hallmarks of cancer (31), it is not likely to be solely sufficient to promote oncogenesis. Very recently, a few studies have explored other tumor-promoting changes that occur in cancers with *ATRX* deficiency, including increased cellular motility in glioma cells and TGF-β activation with *CDH1* (E-cadherin) downregulation in liver cancer cells (16, 32, 33).

Based on its frequent loss in OS and given the important role of *ATRX* in chromatin remodeling and methylation patterns, we sought to define the phenotypic and mechanistic impacts of *ATRX* loss in OS. We hypothesized that loss of *ATRX* would increase aggressive cellular features of OS, including tumor initiation, migration, invasion, and metastasis, and we describe here our investigations into the impact of this *ATRX* loss on OS biology, using a range of models to examine each specific cellular phenotype of aggression. Using RNA-Seq and ATAC-Seq, we examined changes in cellular pathways that correspond with *ATRX* loss. These analyses pinpointed alterations in the NF-κB and several extracellular matrix (ECM)-related pathways. Analysis of chromatin binding motif enrichments identified common overlap of ATRX binding sites with ETS family transcription factor motifs, which is notable because ETS family proteins play an important role in osteogenic differentiation (34, 35). Using high-throughput collateral sensitivity screens, we found that OS cells with *ATRX* knockout were sensitized to an integrin inhibitor. Further examination of these cells demonstrated increased integrin β3 expression with *ATRX* loss. Treatment of *ATRX* knockout cells with the integrin inhibitor was sufficient to reverse the phenotypes of aggression, particularly migration and invasion, and this drug partially reversed the nuclear upregulation of the NF-κB transcription factors. Examining publicly available pan-cancer sequencing data, we similarly found enriched integrin signaling with *ATRX* alteration. The relationship between *ATRX* expression and integrin-binding, NF-κB activation, and ETS family transcription factor binding has not been noted in previous studies, but may impact other known diseases with *ATRX* loss, including other cancers. Our data show that *ATRX* mutations sensitize OS cells to integrin inhibition. Future studies are needed to explore integrin inhibition as a potential new targeted therapy for *ATRX*-deficient OS.

## Results

### *ATRX* loss promotes tumor initiation

We hypothesized that *ATRX* loss-of-expression in OS would correlate with the acquisition of aggressive tumor phenotypes, including alterations in tumor initiation, growth, migration and invasion, and metastasis (Supplemental Figure 2A). We used a range of *in vivo* and *in vitro* models to examine each of these specific phenotypes. To examine how *ATRX* loss alters tumor initiation in OS, we chose to work with a previously-established transgenic *Osterix*-*Cre* mouse model with conditional (floxed) alleles of both *p53* (*p53^fl/fl^*) and *Rb* (*Rb^fl/fl^*) and with a Tet-off cassette providing an additional level of temporal control (36, 37). The transgene expression of Cre recombinase is driven by promoter sequences of *Osterix*, a principal regulator of bone differentiation, and is therefore mostly restricted to committed osteoblast progenitors (36, 38) (although recent studies show *Osterix* expression in additional subsets of cells (39, 40)). These mice with homozygous deletion of *Rb* and *p53* show completely penetrant OS development, typically between 4 and 8 months of age (37), occurring most frequently in the jaw and head, rear limb, hip, ribs, and vertebra. To determine if *ATRX* loss would decrease the time to tumor initiation, we added a floxed *ATRX* allele (*ATRX^fl/fl/y^*) to the *Osx-Cre+p53^fl/fl^Rb^fl/fl^* to create *Osx-Cre+p53^fl/fl^Rb^fl/fl^ATRX^fl/fl/y^*. We removed a doxycycline diet at time of weaning and monitored mice of each genotype for tumor development. To more comprehensively identify tumors that developed in any bone, we performed monthly fluoroscopy on a subcohort of 10 females and 10 males of each genotype to further increase our sensitivity in tumor detection (Supplemental Figure 1A). Tumors were collected and gene recombination of tumors was confirmed by PCR to distinguish the knockout allele from the floxed, non-recombined allele (Supplemental Figure 1B). Consistent with our hypothesis, loss of *ATRX* significantly increased the rate of tumor initiation compared to the *p53*/*Rb* knockout alone (Figure 2A, log-rank p=0.0021).

**Figure 2:**
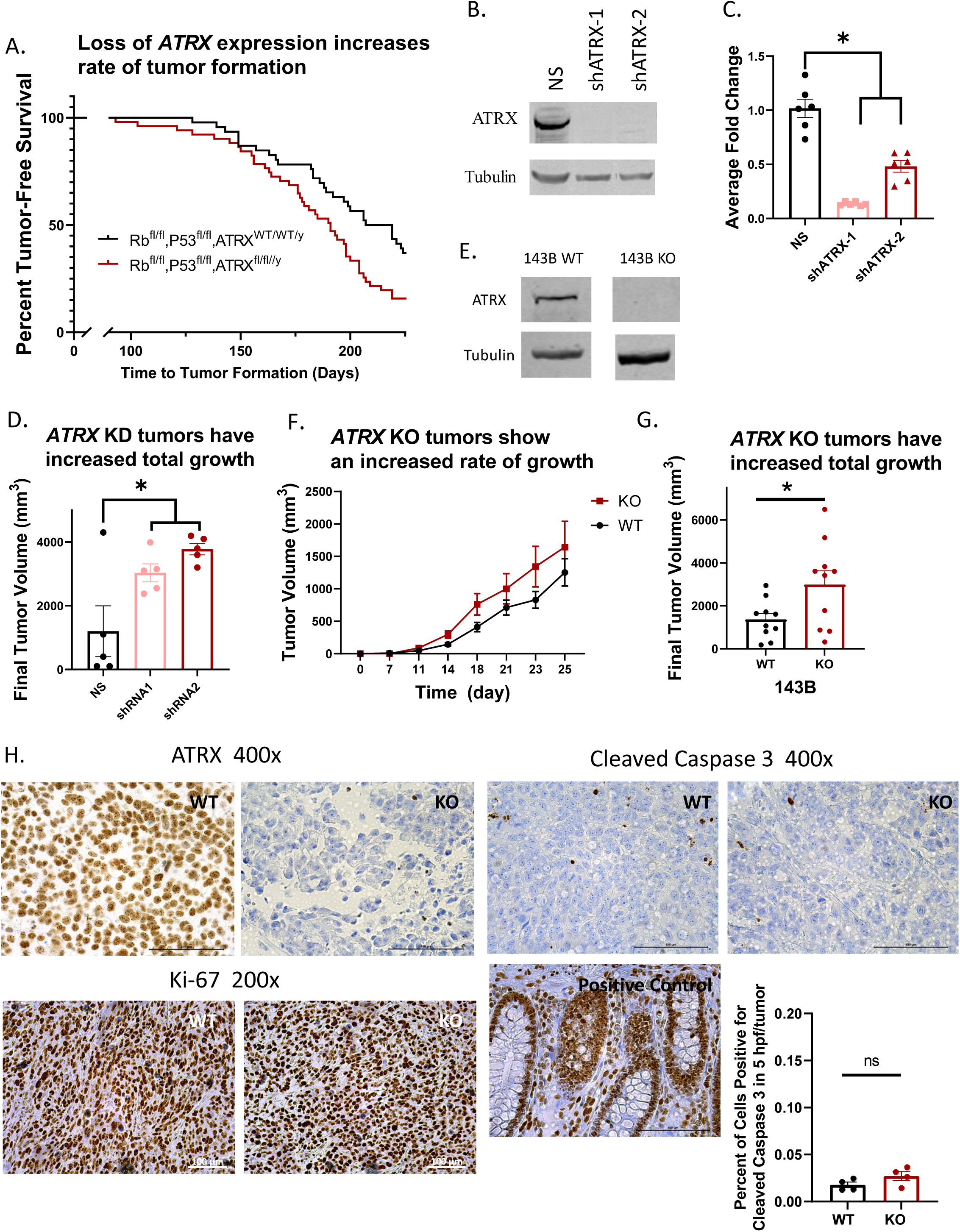
*ATRX* loss promotes tumor initiation and growth. A) Loss of *ATRX* expression increased the rate of tumor formation in an *Osterix-Cre* driven mouse model of OS (Kaplan-Meier, log-rank p=0.0021, experimental cohorts: *Osx-Cre+p53^fl/fl^Rb^fl/fl^* (n=26 males and 25 females) or *Osx-Cre+p53^fl/fl^Rb^fl/fl^ATRX^fl/fl/y^*(n=22 males and 25 females)). B) and C) Western blot and RT-qPCR results showing knockdown (KD) of *ATRX* with two shRNA constructs in the 143B human OS cell line. (p<0.0001 for both NS vs. shATRX-1 and NS vs. shATRX-2 for RT-qPCR results.) D) Established xenografts of the 143B human OS cell line with *ATRX* shRNA knockdown showed greater final tumor volume compared to NS controls (MANOVA, p=0.04 shATRX-1 vs NS, p=0.006 shATRX-2 vs NS, n=3 males and 2 females per treatment group). E) Western blot results show effective CRISPR-Cas9 knockout of *ATRX* expression in the 143B human OS cell line. F) and G) Established xenografts of the 143B cell line with *ATRX* knockout had a faster rate of tumor growth (Repeated measures ANOVA, log transform p=0.02) and larger final tumor volumes compared with wildtype cells (Student’s t-test, p=0.03, n=5 males and 5 females per treatment group). H) Histology of xenograft tumors: The expected *ATRX* expression status of all wildtype and knockout tumors was validated by IHC for *ATRX*. All xenograft tumors stained strongly for Ki-67 (greater than 95% positive) suggesting high cellular proliferation in all tumors. No significant differences were observed in the xenograft tumors with IHC staining for cleaved caspase 3 with less than 5% positively staining cells in all tumors. (Positive control shown.)

### *ATRX* loss promotes tumor growth

Given the role of *ATRX* loss in speeding the time to tumor initiation, we next sought to determine if *ATRX* loss would lead to increased tumor growth. To do this, we chose to compare tumor growth using a xenograft mouse model in which tumors would be easily detected and would develop in the same location (in the subcutaneous flank) for all mice. We stably transduced human 143B OS cells with a non-silencing (GFP) shRNA or one of two independent shRNA constructs targeting *ATRX* (shATRX-1 and shATRX-2). *ATRX* knockdowns were confirmed via Western blotting and RT-qPCR (Figures 2B, 2C). We then injected the control and *ATRX* shRNA knockdown 143B cells subcutaneously in SCID-beige mice and monitored tumor growth rates over time. *ATRX* knockdown significantly enhanced tumor growth in these xenografts for both shRNAs (Figure 2D). To further validate these findings, we also developed a CRISPR-Cas9 knockout (KO) of *ATRX* in the 143B human OS cell line (Figure 2E, Supplemental Figure 2B), and we repeated the previous xenograft experiment with *ATRX* knockout or wildtype cells injected subcutaneously in SCID-beige mice. Consistent with the results of the shRNA-mediated knockdown study, *ATRX* knockout increased both the rate of tumor growth and final tumor volume as compared to wildtype 143B cells (Figures 2F, 2G).

### Histological analysis of xenograft tumors and *in vitro* cells show no significant differences in cell proliferation or apoptosis with *ATRX* loss

In order to investigate whether the increased tumor size found with *ATRX* knockout was due to differences in tumor cell proliferation or apoptosis, immunohistochemistry (IHC) was performed on the formalin-fixed, paraffin-embedded xenograft tumors harvested from these mice. First, we did verify by IHC that the *ATRX* knockout was retained in the final tumors harvested (Figure 2H). Both wildtype and knockout tumors stained very strongly for Ki-67, with greater than 95 percent of cells staining positive, supporting high cellular proliferation in all tumors (Figure 2H). The 143B cell line is known to be an especially proliferative cell line, and perhaps this high baseline of proliferation limits the ability to detect differences in proliferation, if present, between tumors derived from wildtype and *ATRX* knockdown cells.

Additional *in vitro* studies of the 143B non-silenced or shRNA knockdown cells examining change in percent confluence over time in the Incucyte live cell imager showed similarly high proliferation rates of all cells with no significant differences across cell types (Supplemental Figure 2C). Thus, we were unable to conclude that the tumor size differences were due to changes in rate of proliferation. We then performed IHC for cleaved caspase 3 to compare apoptosis, but again, no significant differences were found, with less than five percent of cells for either knockout or wildtype tumors exhibiting positive staining (Figure 2H). Thus, we were also unable to conclude that the tumor size differences were due to resistance to apoptosis.

### *ATRX* loss promotes tumor migration and invasion

We next decided to investigate the impact of *ATRX* loss on changes in cell motility, including migration and invasion, using standard *in vitro* scratch wound and transwell migration/invasion chamber assays. Assays were repeated with two well-established OS cells lines, 143B and MG63, to test for consistency despite the different genetic mutation background found in each cell line. To examine migration, we first performed scratch wound assays with human 143B cells with shRNA-mediated knockdown of *ATRX* and with human MG-63 cells with CRISPR/Cas9-mediated knockout of *ATRX* (Figures 3A-3D, Supplemental Figure 2B). For both cell lines, wound closure rate was significantly increased with *ATRX* knockdown/knockout, supporting increased migration in *ATRX*-null OS cells. We also tested if *ATRX* loss would increase transwell migration/invasion using Boyden chamber migration/invasion assays. As with the scratch wound assays, CRISPR/Cas9-mediated knockout of *ATRX* in both the 143B and MG-63 cell lines increased both migration and Matrigel invasion (Figures 3E-3H).

**Figure 3:**
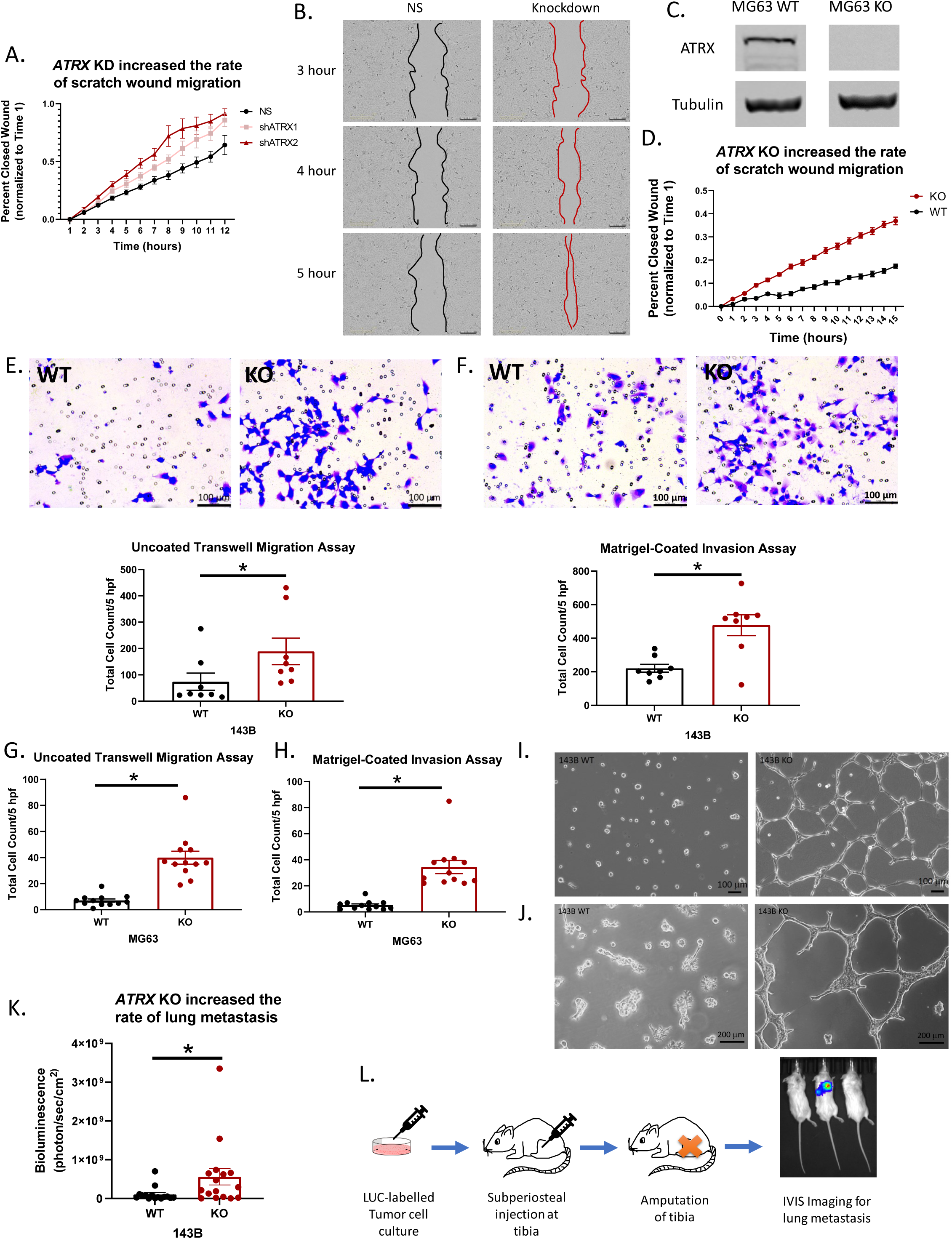
Tumor migration, invasion and rate of metastasis increase with loss of *ATRX* expression. A) In a scratch wound assay, 143B cells with shRNA KD of *ATRX* showed faster wound closure than the non-silenced (NS) control cells (Repeated Measures ANOVA, p<0.0001 for both shATRX-1 and shATRX-2 compared to NS, n=8 replicates per cell type). B) Representative images of scratch wounds at three time points. C) Western blot showing effective CRISPR-Cas9 knockout of *ATRX* expression in the MG-63 human OS cell line. D) Similarly, in the MG-63 cell line, KO cells showed significantly faster wound closure than WT cells (Repeated Measures ANOVA, p<0.0001, WT: n=13 replicates, KO: n=10 replicates). E) and F) 143B KO cells showed increased migration and invasion, respectively, in uncoated and Matrigel-coated transwell plates (Student’s t-test, uncoated p=0.04, n=8 replicates; Matrigel p=0.002, n=8 replicates). G) and H) MG-63 KO cells also showed increased migration and invasion, respectively, in uncoated and Matrigel-coated transwell assays (Student’s t-test, uncoated p<0.0001, n=12 replicates; Matrigel p<0.0001, n=12 replicates). I) After 24 hours of growth on a bed of Matrigel, WT cells remain in tight clusters whereas KO cells form a network of connecting "tubes" or branching networks through the matrix. J) After 96 hours of growth, the distinct differences in morphology between the 143B WT and KO cells are even more apparent. K) Luciferase-labelled 143B WT or KO cells were injected into the subperiosteal space of the tibia of SCID-beige mice. *ATRX* KO correlates with an increased rate of lung metastasis at one week post-amputation (Student’s t-test, p=0.026, WT: n=5 males and 10 females, KO: n=6 males and 11 females). L) Experimental design for orthotopic injections with LUC-labelled cells to study lung metastasis.

Interestingly, when we plated the 143B wildtype and knockout cells on a bed of Matrigel, there was a distinct difference in appearance after 24 hours which was even more apparent after 96 hours. Knockout cells formed a network of connecting “tubes” in the Matrigel, permitting more cell-cell contact, whereas the wildtype cells grew in more confined clusters within the Matrigel (Figures 3I, 3J). These findings support an enhanced ability of knockout cells to invade through the Matrigel, perhaps by secreting enzymes and ECM proteins to form this branching network. When repeating this experiment with the less-proliferative MG-63 cell line, the differences in morphology were less distinct, but still notable after 72 hours (Supplemental Figures 3A, 3B).

### *ATRX* loss promotes tumor metastasis to lungs

The *in vitro* migration and invasion assays suggest *ATRX* loss may promote metastatic dissemination of OS cells. To further examine the role of *ATRX* deficiency in driving metastasis, we used an orthotopic metastasis model of OS. To do this, luciferase-labelled wildtype or *ATRX* knockout cells were injected into the subperiosteal space of the tibia in SCID-beige mice. These mice developed tibial OS in approximately two weeks, after which the affected legs were amputated, and metastatic progression to the lungs was quantified using luciferase imaging with the *In Vivo* Imaging System (Caliper Life Sciences, Inc., PerkinElmer), a method previously validated in our lab as an accurate and accessible assessment of *in vivo* metastatic tumor burden (41). Most of the mice developed lung metastases if given enough time, but consistent with our *in vitro* migration and invasion assays, at one week post-amputation, *ATRX* knockout led to significantly more lung metastases as compared to wildtype cells, supporting the conclusion that *ATRX* loss promotes lung metastasis (3K, 3L).

### *ATRX* loss promotes NF-κB pathway activation and downregulates extracellular matrix (ECM) proteins

Given the important roles of *ATRX* as both a chromatin remodeler and regulator of histone and DNA methylation, we hypothesized that loss of *ATRX* would have broad impacts on gene expression across the genome, and the resulting alterations to multiple cellular pathways would collectively promote more aggressive cancer phenotypes. To examine this, we performed an integrated genomics analysis of RNA-Seq and ATAC-Seq profiles using these non-silenced or shRNA knockdown 143B cells. Analysis of the RNA-Seq data by Gene Set Enrichment Analysis pinpointed enrichment of several pathways relevant to OS upon *ATRX* knockdown, including upregulation of the nuclear factor-κB (NF-κB) pathway and downregulation of ECM proteins (Figures 4A, 4B, Supplemental Figures 4A, 4B). To further understand the mechanisms underlying these gene expression alterations, we analyzed genomic changes in chromatin openness using ATAC-Seq. Analysis of the ATAC-Seq findings revealed altered chromatin openness across the genome upon *ATRX* shRNA knockdown compared to the non-silenced control, supporting the global genomic importance of the role of ATRX as a chromatin remodeler (Supplemental Figure 5A). We then cross-referenced our ATAC-Seq results with our RNA-Seq data and found that transcriptionally upregulated genes were significantly enriched in regions of more open chromatin upon *ATRX* loss (Figure 4C). Similarly, genes that were downregulated with *ATRX* knockdown significantly corresponded with regions of more closed chromatin (Figure 4C). These data suggest that the significant alterations in NF-κB and ECM pathways may derive from *ATRX*-mediated effects on chromatin state. The upregulation of the NF-κB pathway was validated using an ELISA for nuclear extracts of our cell types. Consistent with the RNA-Seq data, NF-κB transcription factor family activity was upregulated in nuclear extracts from both 143B and MG-63 *ATRX* knockout cell lines compared to wildtype cells (Figure 4D).

**Figure 4:**
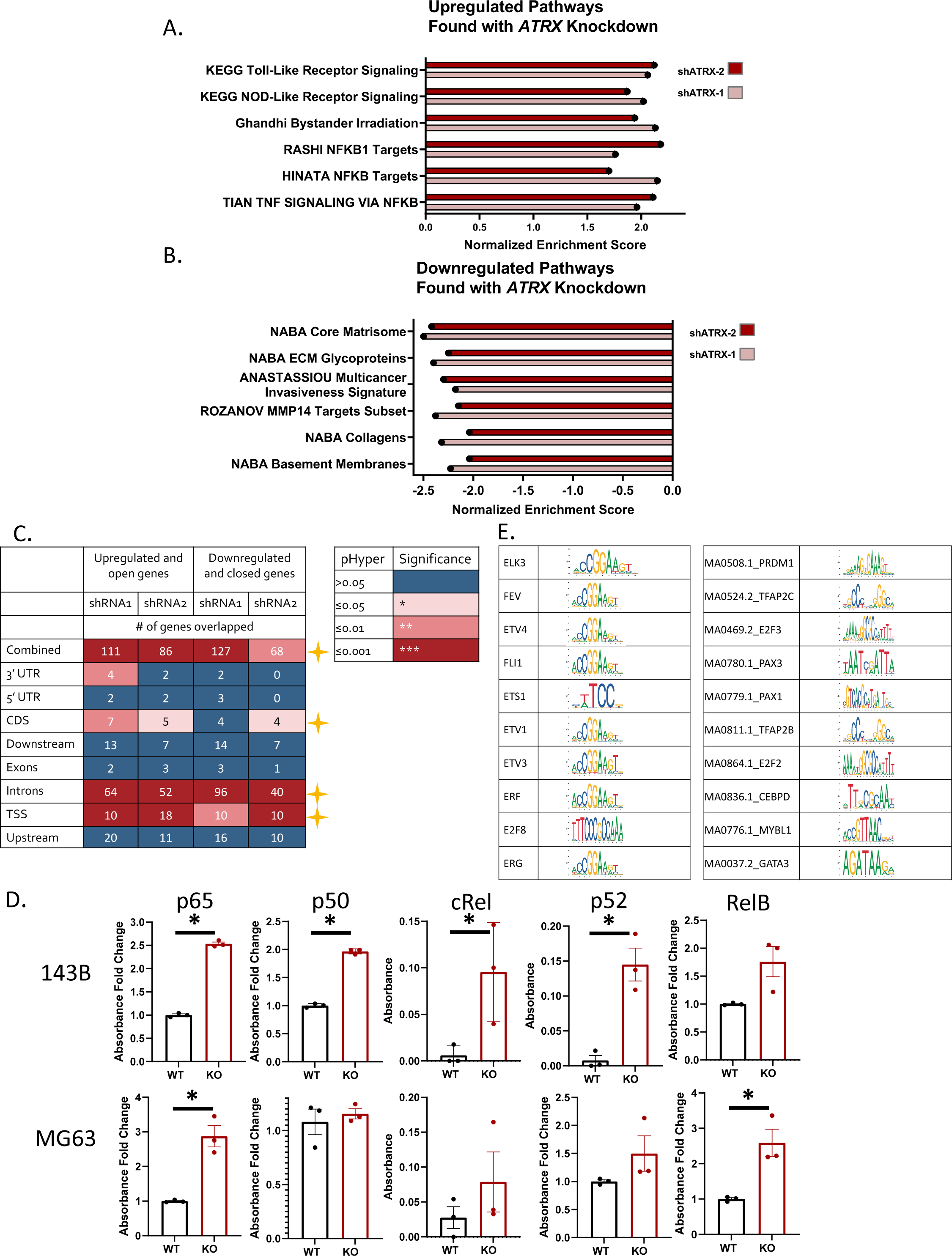
RNA Sequencing identifies upregulation of several NF-κB pathways and downregulation of several extracellular matrix (ECM) pathways. A) and B) RNA-sequencing of 143B human OS cells with either the NS control or one of two shRNA knockdowns of *ATRX* show upregulation of NF-κB pathways and downregulation of various ECM-related pathways with KD of *ATRX* expression. C) ATAC-Seq changes in chromatin openness with *ATRX* shRNA KD correlate with RNA-Seq findings of gene expression changes, particularly when the chromatin peaks are found in regions of introns or transcriptional start sites (Benjamini and Hochberg correction for p value calculations as shown). These data suggest that the significant alterations in NF-κB and ECM pathways may derive from *ATRX*-mediated effects on the chromatin state. D. NF-κB ELISA shows increased relative nuclear expression of this family of transcription factors (TFs) with *ATRX* knockout (KO) when comparing nuclear extracts from both the 143B and MG-63 WT or KO cell lines. (Multiple t-tests; statistically significant comparisons for 143B: p65 p=0.0004, p50 p=0.000009, cRel p=0.04, p52 p=0.009; n=3 replicates for each cell type and TF; statistically significant comparisons for MG-63: p65 p=0.01, RelB p=0.024; n=3 replicates for each cell type and TF). E) Top differentially deviated binding motifs found with *ATRX* shRNA knockdown in the 143B cell line most closely resemble the ETS transcription factor family binding motifs.

### Analysis of common ATRX binding motifs pinpoints ETS family transcription factor binding

To examine potential common binding motifs for *ATRX* across the genome, we analyzed our sequencing data using chromVar to look at known transcription factor binding motifs in the Jasper motif database as well as all 6kmers. The top differentially enriched motifs correspond most closely to ETS family transcription factors, suggesting an important interaction between *ATRX* and these transcription factors (Figure 4E).

### OS cells with *ATRX* loss display collateral sensitivity to pharmacological inhibition of integrin signaling

Our collective data suggest *ATRX* loss provides a selective advantage to OS cells by reducing barriers to tumor initiation, increasing tumor growth rate, increasing migratory/invasive capacity, and enhancing survival in the metastatic niche. Given the host of adaptations and survival benefits conferred on cells with *ATRX* loss, we hypothesized that these cells would be sensitized to loss of a secondary pathway. To identify potential actionable collateral sensitivities, we performed a high-throughput drug screen of 2,100 bioactive compounds on MG-63 wildtype or knockout OS cells. Among the compounds for which this differential viability was outside of the 99^th^ percentile confidence interval, most (74%) displayed increased resistance upon *ATRX* loss (Figures 5A, 5B). These compounds included heat shock protein inhibitors KW-2478, XL888, and VER-49009. Interestingly, however, we also observed increased sensitivity of *ATRX*-null cells to the integrin inhibitor, SB273005 (Figure 5A). SB273005 is a nonpeptide antagonist of the αvβ3 and αvβ5 integrins (42, 43). We further validated this integrin inhibitor sensitization upon *ATRX* loss using IC^50^ assays, confirming a significant drug sensitization with *ATRX* loss (Figure 5C). Integrins are known key interactors with ECM components and directly activate the NF-κB pathway, all of which is consistent with our observations that both NF-κB and ECM pathways are altered upon *ATRX* loss.

**Figure 5:**
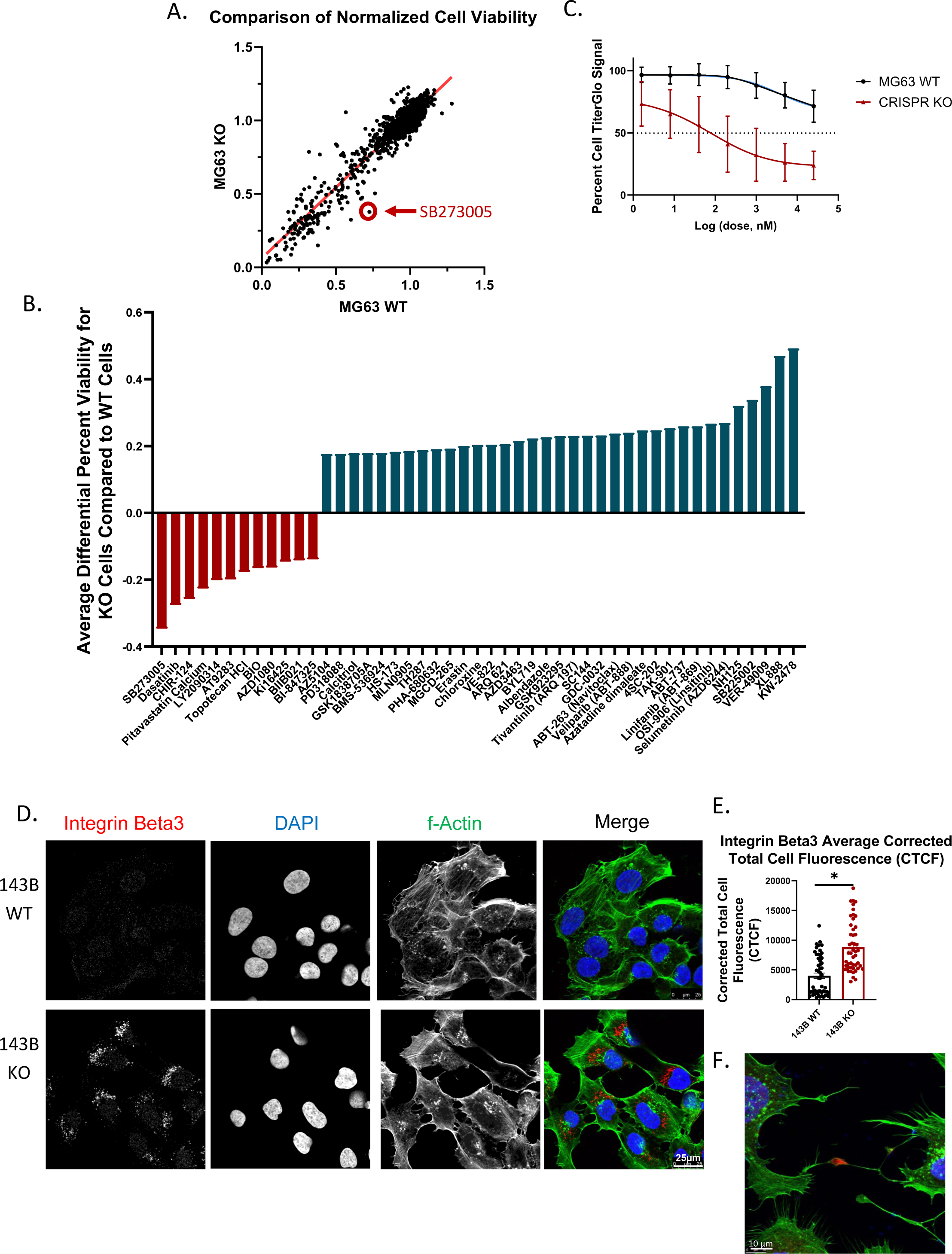
*ATRX* KO sensitizes cells to pharmacological inhibition of integrin signaling *in vitro* and KO cells have a greater expression of integrin β3. A) Comparison of normalized cell viability for all 2100 drugs in a bioactive compound screen. *ATRX* KO cells were most significantly sensitized to the integrin inhibitor SB273005. B) Among the compounds for which this differential viability was outside of the 99th percentile confidence interval, most (74%) displayed increased resistance upon *ATRX* loss. C) SB273005 IC^50^ curves show significant sensitization of KO cells to the integrin inhibitor SB273005. D) and E) 143B *ATRX* KO cells show significantly increased expression of integrin β3 compared to WT cells (Student’s t-test, p<0.0001, KO n=49, WT n=52). F) Example of integrin β3 expression at cell-cell adhesions.

The integrin inhibitor, SB273005, has a high affinity for αvβ3 integrins, and so we hypothesized that there would be an increase in expression of integrin αvβ3 at the cell surface with loss of *ATRX* expression. To test this, we performed immunofluorescent imaging of our cell lines. As predicted, we saw significantly increased expression of integrin β3 in our knockout cells in both the 143B and MG-63 cell lines (Figures 5D-5F, Supplemental Figure 3C). One of the key matrix components to which integrins αvβ3 and αvβ5 bind is osteopontin, and so we examined expression of this gene. Osteopontin mRNA expression was upregulated in our *ATRX* knockdown cells (Supplemental Figure 3D), and we hypothesize that there may be increased secretion of this phosphoprotein correlated with *ATRX* loss. Future studies will further investigate these ECM-integrin relationships and how they correlate with *ATRX* deficiency.

We next tested the integrin inhibitor SB273005 *in vivo* with xenograft mouse tumors to examine its efficacy as a therapeutic for *ATRX*-deficient OS. For this experiment, we chose to use an established *ATRX*-null OS cell line, U-2 OS, to see if this cell line would respond *in vivo* as the MG63 knockout cells did *in vitro*. As observed in our high-throughput *in vitro* screen, treatment with an integrin inhibitor, SB273005, significantly reduced *in vivo* tumor growth in xenografts formed from subcutaneous flank injections of these U-2 OS cells (Figures 6A, 6B).

**Figure 6:**
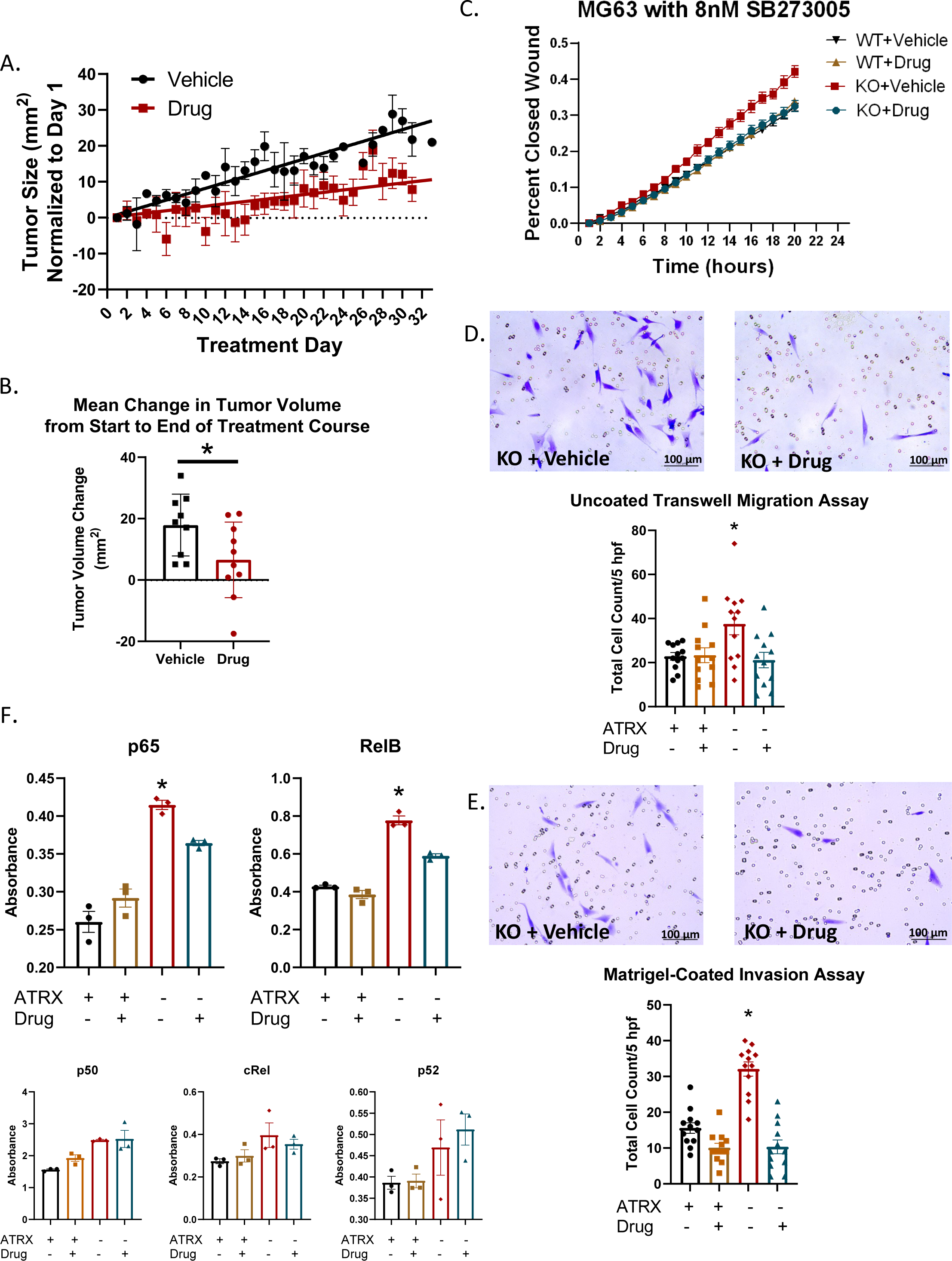
*ATRX* KO sensitizes cells to pharmacological inhibition of integrin signaling *in vivo*, and integrin inhibition is sufficient to reverse aggressive phenotypes seen with *ATRX* loss. A) and B) Treatment with the integrin inhibitor, SB273005, significantly reduced tumor growth in *ATRX*-null U-2OS cells (Repeated Measures ANOVA, p<0.0001 for volume change over time; Student’s t-test, p= 0.02 for final tumor volume change; vehicle-treated: n=4 females, 5 males; integrin inhibitor-treated: n=5 females, 5 males). C) The integrin inhibitor reversed the increased rate of migration conferred by loss of *ATRX* in the MG-63 cell line (Repeated Measures ANOVA, p<0.0001, WT Vehicle n=19 replicates; WT Drug n=26 replicates, KO Vehicle n=18 replicates, KO Drug n=25 replicates). D) and E) The integrin inhibitor, SB273005, reversed the increased migration and invasion observed with the MG-63 *ATRX* KO cells in both uncoated transwell assays (D) and Matrigel-coated transwell assays (E). (Tukey’s multiple comparisons test for uncoated wells: *ATRX* KO Vehicle vs. WT Vehicle p= 0.028, KO Vehicle vs. WT Drug p= 0.035, KO Vehicle vs KO Drug p= 0.0117, n=12 wells. Tukey’s multiple comparisons test for Matrigel-coated wells: *ATRX* KO Vehicle vs. WT Vehicle p<0.0001, KO Vehicle vs. WT Drug p<0.0001, KO Vehicle vs KO Drug p<0.0001; n=12 wells). F) Integrin inhibitor treatment partially reversed the upregulation of NF-κB transcription factors p65 and RelB in the *ATRX* KO cells. (Tukey’s multiple comparisons tests for p65: ATRX KO Vehicle vs. WT Vehicle p<0.0001, KO Vehicle vs. WT Drug p<0.0001, KO Vehicle vs KO drug p=0.028; n=3. Tukey’s multiple comparisons tests for RelB: *ATRX* KO Vehicle vs. WT Vehicle p<0.0001, KO Vehicle vs. WT Drug p<0.0001, KO Vehicle vs KO drug p=0.0002; n=3.)

### Integrin inhibition is sufficient to reverse aggressive phenotypes seen with ATRX loss

Given the functional connection between αvβ3 and αvβ5 signaling, NF-κB signaling, and invasive phenotypes in cancer, we tested if integrin signaling inhibition could reverse the increase in phenotypes of aggression that we found with *ATRX* knockdown/knockout. We first repeated our scratch wound assays with wildtype and knockout cells treated with vehicle control or with 8 nM SB273005. The knockout cells treated with vehicle still retained a significant increase in rate of wound closure relative to both vehicle- and drug-treated wildtype cells, however the knockout cells treated with SB273005 had a wound closure rate that was very similar to the wildtype cells (Figure 6C, Supplemental Figure 3E). Thus, this experiment supports that the integrin inhibitor is sufficient for reversal of the increased migratory phenotype conferred by loss of *ATRX.* These results suggest that the enhanced migratory capability of *ATRX*-deficient cells is likely due to increased integrin expression and binding. Similarly, treatment of knockout cells with the integrin inhibitor reversed the increased migration and invasion observed in the transwell migration and Boyden invasion chamber assays (Figure 6D, 6E).

### Integrin inhibition partially reverses upregulation of NF-κB signaling

Given the reported relationship between integrin binding and NF-κB signaling, we also investigated whether the integrin inhibitor would be sufficient to reverse the upregulation of the NF-κB transcription factor family in the *ATRX* knockout cells. Both knockout and wildtype MG-63 cells were incubated for 24 hours with either SB273005 or the vehicle control, nuclear extractions were performed, and the NF-κB ELISA was repeated. The two significantly upregulated transcription factors seen in our prior experiment with the MG-63 knockout cells, p65 and RelB, were both significantly rescued by integrin inhibitor treatment compared to vehicle control (Figure 6F). These results support a close correlation between the increased migratory and invasive phenotypes found with *ATRX* deficiency, integrin binding and NF-κB pathway activation.

### Integrin signaling is enriched in *ATRX*-altered tumors in ICGC/TCGA Pan-Cancer Analysis of Whole Genomes dataset

In order to explore whether similar cellular signaling alterations are found in human cancers with altered *ATRX* expression, we used cBioportal to examine the gene expression data of the ICGC/TCGA Pan-Cancer Analysis of Whole Genomes dataset (18, 19, 44). Importantly, in line with our own experimental findings, we did find integrin signaling to be enriched in the *ATRX*-altered subset of tumors (Figure 7A, 7B). Additionally, survival in the subset of patients with *ATRX*-altered tumors was significantly decreased compared to those with *ATRX* wild-type expression (Figure 7C). These results further support a close correlation between *ATRX* deficiency, survival, and integrin signaling.

**Figure 7:**
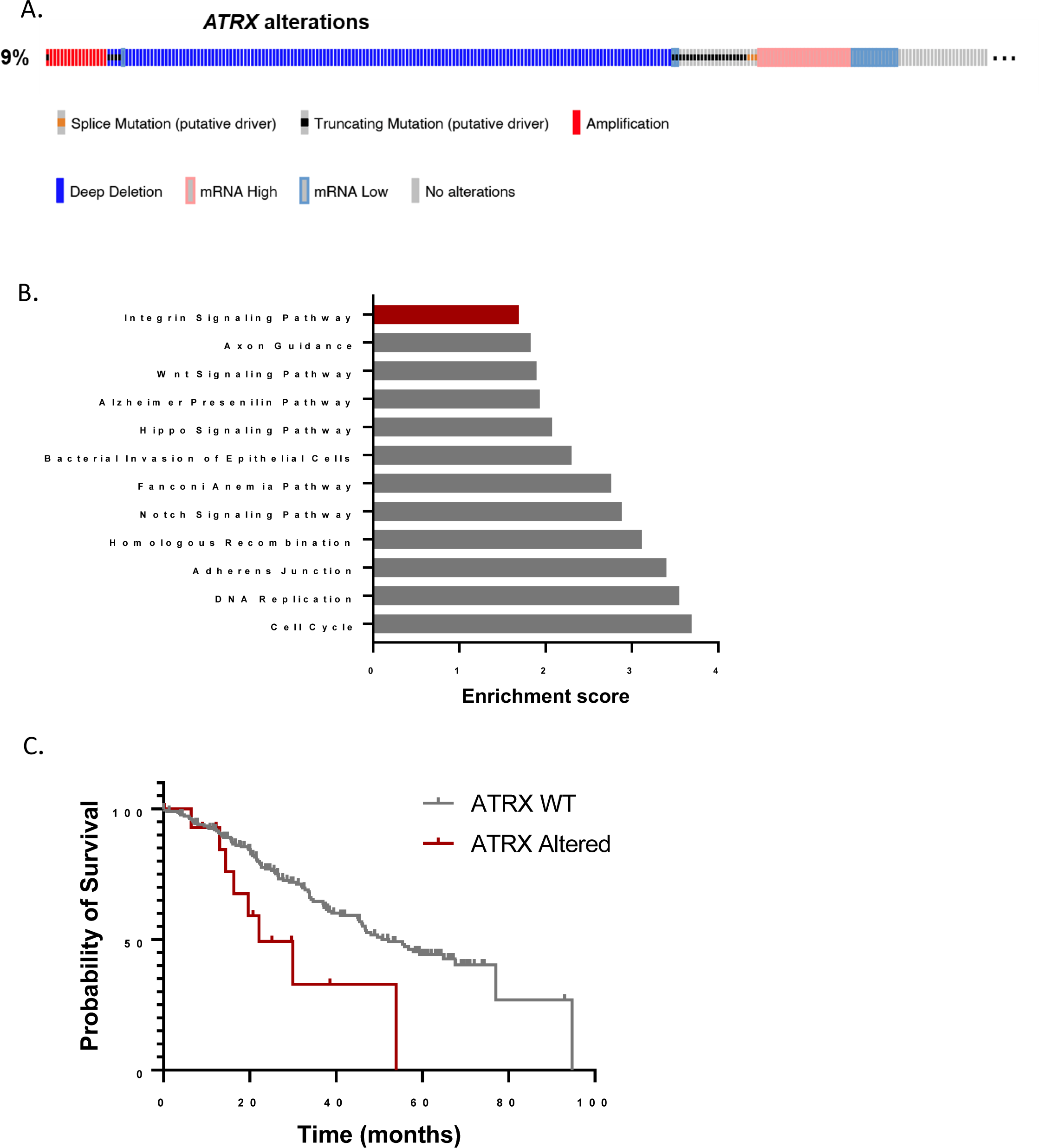
Integrin signaling is enriched in *ATRX*-altered tumors in the ICGC/TCGA Pan-Cancer Analysis of Whole Genomes sequencing dataset. A) Oncoprint of *ATRX* status. *ATRX* was found to be altered in 9% of 2583 samples after “hiding mutations and copy number alterations of unknown significance” in the ICGC/TCGA Pan-Cancer Analysis of Whole Genomes dataset on cBioPortal (18, 19, 44). B) Integrin signaling was enriched in the *ATRX*-altered tumors. Over-representation Analysis (ORA) was performed on expression data between *ATRX*-altered and unaltered groups (105). Genes submitted to ORA had higher expression in *ATRX*-altered groups than unaltered groups, and all identified genes had q values < 0.05. The query was submitted to the Panther, KEGG, and Wikipathway cancer databases. Enrichment ratios for identified pathways with FDR < 0.05 are displayed. C) Kaplan-Meier survival curve based on *ATRX* status. Hazard ratio = 3.917, 95% CI is 1.335 – 11.5, p<0.036. Note, not all *ATRX*-altered tumors have deletions—aside from homdel, there are two missense mutations and one amplification represented in the altered group.

## Discussion

Our experimental results support the hypothesis that *ATRX* loss, common in human OS, plays a key role in enhancing the aggressiveness of OS. Loss of expression of this gene increases multiple oncogenic phenotypes, including increased tumor initiation, growth, migration, invasion, and metastasis. Our investigations into the underlying cellular mechanisms driving these aggressive OS phenotypes with *ATRX* deficiency point to changes in the NF-κB pathway, ECM protein expression, ETS transcription factor binding, and integrin expression, specifically integrins αvβ3 and αvβ5. *ATRX*-deficient cells display substantially increased sensitivity to integrin signaling inhibition. Additionally, examination of pan-cancer sequencing data supports these correlations between *ATRX* deficiency, survival, and integrin signaling across a range of cancer types.

NF-κB activity is known to increase tumor cell proliferation, suppress apoptosis, and promote angiogenesis (45, 46). NF-κB signaling also enhances tumor invasiveness by inducing and maintaining epithelial-mesenchymal transitions required for tumor metastasis (47). Specifically in OS, Felx et al. (48) found that NF-κB pathway activation played a central role in proliferation in the MG-63 human OS cell line, and Zhao et al. (49) reported that the NF-κB pathway was a key regulator of OS tumor growth, metastasis and resistance to chemotherapeutics. We demonstrated increased expression of integrin β3 with *ATRX* knockout in our cells. Scatena et al. (50) linked NF-κB activation with integrin αvβ3 binding to its ECM ligands. ECM components include various proteins and growth factors, such as osteopontin, fibronectin, collagens, proteoglycans, and laminins. OS cells adhere to the matrix via cell-surface receptors, primarily integrins, which bind to these ECM proteins (51). The ECM plays a critical role in tumor migration and metastasis as tumor cells use integrin binding as well as various ECM-degrading proteases, including MMPs, to invade and metastasize (52). Several studies have examined the general interplay between these cellular pathways and OS biology. Li et al. (53) were able to inhibit OS metastasis with the combined blockade of both NF-κB signaling and integrin-β1 expression in MG-63 cells. Very recently, Shi et al. (54) showed that expression of avβ3 integrins and fibronectin were both correlated with poor clinical prognosis and decreased survival in OS patients. These findings all point to a precise mechanism through which OS with *ATRX* loss behave more aggressively in the clinical setting.

Our successful attenuation of xenograft tumor growth as well as tumor cell migratory and invasive capabilities with the integrin inhibitor SB273005 supports the importance of integrin binding for increased OS aggression correlated with *ATRX* loss. In ecological contexts, it is often the case that an advantage in one environment leads to a ‘*collateral sensitivity’* to another environment. Given the host of adaptations and survival benefits conferred on cells with *ATRX* loss, we hypothesized that these cells would harbor some collateral sensitivity to loss of a secondary pathway. SB273005 is a potent, orally active nonpeptide integrin inhibitor with a high affinity for integrin αvβ3 (Ki=1.2 nmol/l) and a somewhat lower affinity for integrin αvβ5 (Ki=0.3 nmol/l) (43). Research on this inhibitor is somewhat limited to date but has included studies of its effect on bone resorption and osteoporosis, arthritis, and the production of Th2 cells and cytokine IL-10 in pregnant mice (42, 43, 55). Gomes et al. (56) studied breast adenocarcinoma cells in whole blood under flow conditions and found that a combination of SB273005 and lamifiban (a non-peptide antagonist specific for platelet αIIbβ3) successfully inhibited adhesion to the vascular ECM. Several other αvβ3-targeting drugs have advanced to clinical trials for treatment of various solid tumors, including cilengitide, etaracizumab, and the small molecule GLPG0187 (57–62). Despite success in early clinical trials, many of these therapies did not produce clinical outcomes significantly improved compared to standard treatment regimens. Based on our own findings, one might ask whether a more defined target population, based on specific cellular signaling pathway alterations and gene mutations, such as *ATRX* loss, would improve these outcomes. Indeed, such a similar dependence on specific mutations for drug efficacy has been demonstrated with the use of PARP inhibitors most successfully in breast cancer tumors harboring mutations in the *BRCA1* and *BRCA2* genes (63).

In addition to altered integrin expression, we discovered increased mRNA expression of osteopontin following *ATRX* loss. Osteopontin (*OPN* or *SPP1*) is an important component of the ECM in bone, and high expression and secretion of osteopontin in numerous cancer types, including breast, prostate, lung, gastrointestinal, hepatocellular, cervical, and bladder cancers, have been clinically correlated with poor prognosis and shortened survival times (64–73).

Gaumann et al. (74) also showed that strong expression of osteopontin correlated with progression of malignancy and metastasis in poorly differentiated sarcomas. Song et al. (75) showed that targeting osteopontin expression with miR-4262 could reduce cell invasion and migration in OS cells. In line with our study findings, binding of osteopontin to integrin αvβ3 directly activates the NF-κB pathway (50). Future assays will examine changes in the secreted proteins, including osteopontin, to further characterize how *ATRX* expression levels affect these ECM proteins.

The examination of *ATRX* binding motifs from our sequencing data shows a close similarity to ETS family transcription factor binding motifs. ETS family proteins are also involved in osteogenic differentiation and have been reported to play an important role in osteoblast development and bone formation (34, 35). Studies show that the ETS family of transcription factors plays a crucial role in driving malignancy of tumor cells by prevention of apoptosis, support of angiogenesis, and promotion of invasion and metastasis (34, 35, 76–82). ETS factors are well-known critical mediators of ECM remodeling and invasive properties in cancers, regulating a wide spectrum of ECM-related target genes (34, 83). Also fitting with our own experimental findings, ETS transcription factors are known to share crosstalk with NF-κB signaling, and ETS-1 and ETS-2 have been shown to bind directly to the promoter regions of integrins αv and β3 as well as their ligand, osteopontin (77, 81, 84–88).

Our research has not completely elucidated how ATRX expression alters ETS transcription factor activity, but the enrichment of the ETS binding motifs with changes in chromatin accessibility suggest that functional ATRX may repress these transcription factors by maintaining closed chromatin, effectively preventing the tumor-promoting sequelae that would otherwise occur with active ETS protein binding. Intriguingly, Li et al. (89) demonstrated that ETS-1 and Daxx protein colocalize to the PML (pro-myelocytic leukemia) nuclear bodies. DAXX/EAP1 (Ets1-associated protein 1) is able to bind to the N-terminal part of ETS-1 and cause repression of transcriptional activation of ETS-1 target genes, including *MMP1* and *BCL2* genes (89). In more recent studies, researchers have found that ATRX also localizes to the PML nuclear bodies and forms a binding complex with Daxx at this location (90, 91). Future studies should explore whether ATRX also binds with ETS-1 in PML bodies, similar to DAXX/EAP1, and in this way, directly represses transcriptional activation of ETS-1 target genes.

As we have described, the interactions between ETS binding, NF-κB activation, integrin αvβ3 expression, and osteopontin expression have been closely linked with more aggressive tumor biology in prior studies across cancer types, but the association with *ATRX* expression has not been noted previously. This relationship may pertain to the underlying cellular signaling pathways and pathogenesis of other *ATRX*-deficient diseases, including other cancers and ATR-X mental retardation syndrome. In a large pan-cancer sequencing dataset, integrin pathway signaling was enriched in this *ATRX*-mutated population, supporting the hypothesis that our findings will apply to a wide range of *ATRX*-deficient cancers. Our graphical model of these signaling pathways based upon our experimental findings is shown (Figure 8). Understanding how *ATRX* alters these cellular pathways in OS may aid in identifying new targeted therapeutics for *ATRX*-deficient OS tumors, including the potential use of current and novel integrin inhibitors. Future research should investigate further the role of ECM alterations in OS metastasis and the efficacy of integrin inhibitors as a targeted therapy for specific subsets of OS. How *ATRX* loss alters these pathways in other cancers should also be explored.

**Figure 8:**
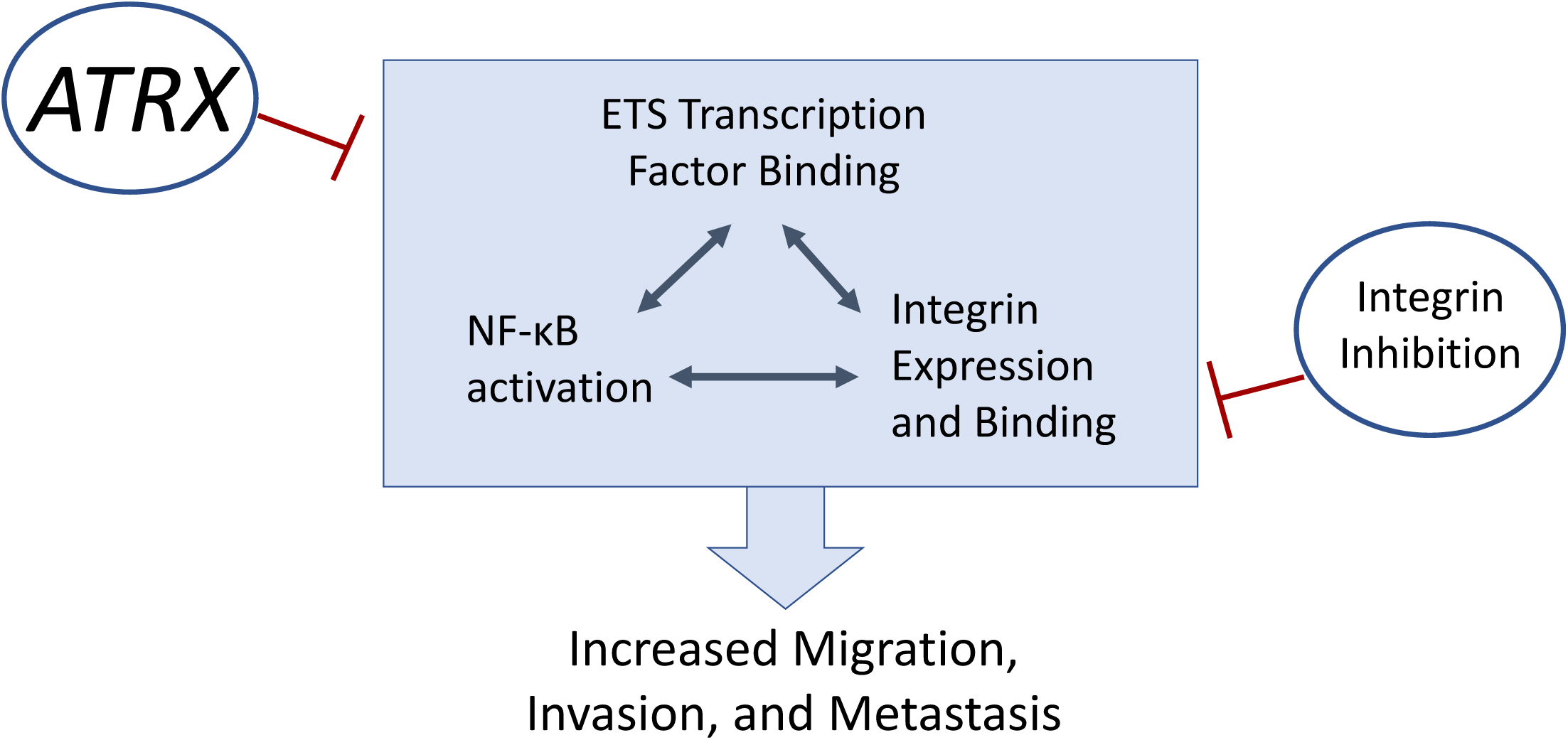
Graphical working model. Numerous studies have shown a close interaction between ETS family transcription factor binding, NF-κB pathway activation, and integrin expression and binding, all of which collectively promote increased cellular motility and metastasis in cancers. Our study has now revealed that *ATRX* expression suppresses these pathways to prevent the oncogenic phenotypes of migration, invasion and metastasis, and integrin inhibition may be a potential new targeted therapy for *ATRX*-deficient OS.

## Materials and Methods

### Cell Culture

143B and MG-63 human osteosarcoma cell lines as well has HEK293T cells were cultured in Dulbecco’s modified Eagle’s medium (Gibco; catalog #11995065) supplemented with 10% fetal bovine serum (HyClone; catalog #SH30071.03HI) and 1% penicillin-streptomycin (Gibco; catalog #15140122). The U-2 OS human osteosarcoma cell line was cultured in McCoy’s 5A modified medium (Gibco; catalog #166000082) supplemented with 10% fetal bovine serum (HyClone; catalog #SH30071.03HI) and 1% penicillin-streptomycin (Gibco; catalog #15140122). All cell lines were incubated at 37°C and 5% CO2 in a humidified incubator. All cell lines were obtained from the Duke University Cell Culture Facility, which performs routine *Mycoplasma* testing and verifies cell identity by analysis of short tandem repeats. Cell line authentication was verified again after *ATRX* knockouts were created, and chromatographs are provided (Supplemental Figure 6). 143B and U-2 OS cells are from a female patient, and MG-63 cells are from a male patient.

### Western blotting

Total protein was extracted from cultured cells by washing cells twice in chilled 1x Dulbecco’s phosphate-buffered saline (DPBS) (Gibco; catalog #14190144) and lysing them in 1x radioimmunoprecipitation assay (RIPA) buffer (Thermo Fisher; catalog #89900) supplemented with 2x Halt protease inhibitor cocktail (Thermo Fisher; catalog #78440). Cell lysates were incubated at 4°C for 15 minutes with rocking and clarified by centrifugation at high speed in a benchtop centrifuge for 10 minutes, after which clarified lysates were incubated for 5 minutes at 95°C in 1x Laemmli gel loading buffer (Thermo Fisher; catalog #AM8546G). Lysates were separated in 4 to 12% NuPAGE Novex Bis-Tris gels (Thermo Fisher; catalog#NP0321 or #NP0323) while in 1x NuPAGE MOPS SDS running buffer (Thermo Fisher; catalog #NP0001). Proteins were transferred onto nitrocellulose membranes (GE Healthcare; catalog #10600012) in 1x NuPAGE transfer buffer (Thermo Fisher; catalog #NP0006) at 50 V for 2 hours at 4°C. Membranes were blocked for 1 hour at room temperature or overnight at 4°C in blocking buffer (2.5 g of dried milk powder in 50 mL of TBST; TBST was made with 100 mL of 10x TBS, 10 mL of 10% Tween, with deionized water added to bring to 1L volume). Membranes were then washed three times for 5 minutes with 1x DPBS (Gibco; catalog #14190144) with 0.05% Tween 20 (PBS-T). Primary antibodies (Anti-ATRX: Cell Signaling Technology #14820; Anti-α-tubulin: Abcam #ab7291) were diluted in blocking buffer (1:1000 for ATRX, and 1:5000 for α-Tubulin). These antibodies were added to the membranes, incubated for 1 hour at room temperature or overnight at 4°C, washed three times for 5 minutes in 1x DPBS (Gibco; catalog #14190144) with 0.05% Tween 20 (PBS-T), and the incubated for 1 hour at room temperature with IRDye-coupled goat anti-rabbit 800CW (LI-COR; catalog #925-32211) or goat anti-mouse 680RD (LI-COR; catalog #925-68070) secondary antibodies diluted 1:20,000 in blocking buffer. Images were captured on an Odyssey Fc imager (LI-COR).

### RT-qPCR

RNA was extracted from cultured cells following the protocol of the ReliaPrep RNA Cell Miniprep System (Promega; catalog #Z6012). Concentrations of the RNA extractions were measured with a Thermo Fisher NanoDrop One spectrophotometer, and all RNA samples were diluted to equal the sample with the lowest concentration. The Applied Biosystems High-Capacity cDNA Reverse Transcription kit (Applied Biosystems; catalog #4374966) was used for reverse transcription. Samples were incubated in a thermocyler at 25°C for 10 minutes, 37°C for 2 hours, 85°C for 5 minutes, and then brought to 4°C. Final cDNA samples were diluted 1:5 with molecular grade water (Invitrogen; catalog #10977015). qPCR was performed in Applied Biosystems MicroAmp Fast Optical 96-well barcoded reaction plates (Thermo Fisher; catalog #4346907). Samples were added to 10 uM forward and reverse primers and 2x KAPA SYBR Fast Universal qPCR Master Mix (Kapa Biosystems; catalog #KK4602). Primers for the housekeeping gene, HPRT1, were as follows: forward 5’-GAA AAG GAC CCC ACG AAG TGT-3’ and reverse 5’-AGT CAA GGG CAT ATC CTA CAA-3’. Primers for ATRX were as follows: forward 5’-TCC TTG CAC ACT CAT CAG AAG AAT C-3’ and reverse 5’-CGT GAC GAT CCT GAA GAC TTG G-3’.

### shRNA knockdown

The ATRX shRNA clones TRCN0000013588, TRCN0000013589, TRCN0000013590, TRCN0000013591, and TRCN0000013592 were obtained from Sigma Mission shRNA collection (https://www.sigmaaldrich.com/life-science/functional-genomics-and-rnai/shrna/individual-genes.html). A non-silencing PLKO-GFP plasmid also was obtained from the same Sigma Mission shRNA collection (catalog #SHC005). HEK293T cells were plated at 500,000 cells/well of a 6-well culture plate. Twenty-four hours later, for each well of cells, lentiviral packaging plasmids (1.8 µg pΔ8.9 and 0.2 µg pCMV-VSV-G (Addgene; catalog#8454)) were combined with 2 µg shRNA plasmid in 200 µL Opti-Mem (Thermo Fisher; catalog #11058021) in a centrifuge tube. In a corresponding centrifuge tube, 4 µL Lipofectamine 2000 (Invitrogen; catalog #11668019) was added to 200 µL Opti-Mem. The plasmid mixture was added to the Lipofectamine mixture and incubated for 20 minutes at room temperature. The wells of HEK293T cells were washed with 1x DPBS (Gibco; catalog #14190144) and then 800 µL Opti-Mem was added to each well. The 400 µL mixture of plasmids and Lipofectamine were then added to each well. Plates were incubated at 37°C and 5% CO2 in a humidified incubator for 3 hours. Then, the wells were aspirated and washed with 1 mL DPBS before adding 2 mls of Dulbecco’s modified Eagle’s medium (Gibco; catalog #11995065) supplemented with 10% fetal bovine serum (HyClone; catalog #SH30071.03HI) and 1% penicillin-streptomycin (Gibco; catalog #15140122), and plates were returned to the incubator. Target 143B cells were plated 24 hours later at 500,000 cells/well of a 6-well culture plate, and after another 24 hours, viral media was removed from the HEK293T cells with a 10 mL syringe, filtered through a 0.45 μm Polyethersulfone Membrane (VWR; catalog #28145-505) into a 15 mL conical tube containing 4 µL Polybrene. The HEK293T cells received fresh complete media, and each target 143B well was aspirated before adding the viral media and continuing the incubation. This viral media transfer was repeated 24 hours later, and another 24 hours later, cells were passaged and moved into 10 cm round tissue culture dishes. 3 µL of puromycin at 2 mg/mL was added to each culture dish to select for cells with successful shRNA plasmid uptake. Each plasmid was allowed to multiply to confluence in a T75 plate, and then clones were tested for *ATRX* expression by both qPCR and Western blotting to confirm *ATRX* knockdown. The two strongest knockdowns were used with the non-silenced control for subsequent experiments.

### CRISPR-Cas9 knockout of *ATRX*

sgRNA sequences targeting exon 4 of *ATRX* were designed using ChopChop (92) and CasOffinder (93) and cloned into the px459 vector from Feng Zhang, for coexpression of a sgRNA with S. pyogenes Cas9 and a puromycin resistance marker (Addgene plasmid #62988). The sgRNA sequences used for generating the knockout cell lines are CAGGATCGTCACGATCAAAG (ATRX-3) and TCGTGACGATCCTGAAGACT (ATRX-4) and are designed to generate a 20 bp deletion when used together. MG-63 or 143B knockout cells were created by cotransfection of ATRX-3 and ATRX-4 sgRNA vectors using TransIT-LT1 (Mirus) according to manufacturer’s suggestions, followed by transient selection with puromycin. Surviving cells were expanded and re-plated into 96 well plates at limiting dilution to isolate clonal lines. Clones were screened for CRISPR-mediated deletion by loss of a Tsp45I restriction site and positive clones were Sanger sequenced to confirm and characterize the genomic deletion event. Successful knockout of *ATRX* was also verified by Western blot and RT-qPCR.

### Luciferase labelling of cells

143B WT and *ATRX* CRISPR-Cas9 knockout cells were transfected with the pLenti CMV Puro LUC plasmid (Addgene #17477) following the same transfection protocol as for the shRNA plasmids described above. Following puromycin selection of all transfected cells, each LUC-labelled cell line was passaged and plated at 500 cells/plate in 10-cm tissue culture plates. After about one week, single colonies of cells were harvested by gently scraping the individual clone’s colony in the plate with a 200 uL pipet tip and then transferring to a single well of a 24-well cell culture plate. Multiple colonies were collected in this way and then each isolated clone was allowed to expand to confluence. The Dual-Glo Luciferase Assay (Promega; catalog #E2920) was then performed and luciferase fluorescence levels were read on the SpectraMax plate reader to identify individual WT and KO clones that expressed equal levels of luciferase (Supplemental Figure 2D). The two most similar clones were then expanded for use in the orthotopic injections described below.

### Growth of cells on Matrigel bed

100 μl of Matrigel (Corning, 354230) was evenly spread in each well of a pre-chilled 24-well plate. The plate was placed in an incubator for 15 minutes to allow the Matrigel to set. Then, 5 x 10^3^ cells per well were seeded onto the Matrigel bed. Cells were imaged every 24 hours using the ZEISS AxioVert microscope.

### In vitro immunofluorescent staining of cells

Coverslips (Electron Microscopy Cat. #72230-01) stored in 90% EtOH were placed on parafilm in a petri dish and washed 3x with sterile 1X PBS. Coverslips were coated with 10ug/mL human fibronectin for 1hr at 37°C then gently washed 3x with 1XPBS. Coverslips were then carefully transferred to 24-well plate (with #5 watchmaker’s tweezers), and cells were plated at ≤50% cell density and allowed to spread for 12 to 24 hours. Media was removed and coverslips were gently washed once with 1XPBS at room temperature. Cells were fixed with cold 4% PFA for 10 minutes then washed 2-3x with 1XPBS. Coverslips were moved to staining chamber (p1000 tip box with parafilm and lid) and 1XPBS was quickly added. To permeabilize, PBS was aspirated off and 0.1% Triton-X-100 in PBS was added for 5 minutes at room temperature. Coverslips were then washed 3x 5 minutes with 200ul 1XPBS. Coverslips were blocked with 5% BSA for 30minutes. Primary antibodies (Alexa Fluor 488 Phalloidin, Cell Signaling, cat#8788; Integrin β3, Thermo Fisher, cat#13166) were diluted 1:200 in 1% BSA in PBS, and cells were stained either 1 hour at room temperature or overnight at 4°C in humidified chamber. Antibodies were aspirated and coverslips were washed 3x 5minutes with 1XPBS. Secondary antibodies (Thermo Fisher, cat#A10523) were added at 1:1000 in 1% BSA for 1 hour at room temperature. Secondary antibodies were aspirated and coverslips were washed 3x 5minutes with 1XPBS. Coverslips were gently inverted, with excess PBS removed, mounted onto slides with mounting media containing DAPI (Thermo Fisher, cat#P36931) and sealed with nail polish. Imaging was performed on a Leica SP5 Confocal Laser Scanning Microscope. Images were captured with a 40X oil objective. Images were analyzed using ImageJ (94).

### Scratch wound assays

143B NS and shATRX cells were plated in separate wells of a 96-well culture plates at 2 x 10^4^ cells/well in 200 uL of culture media. After 24 hours, cells had reached confluence. A sterile 20 uL pipet tip was manually scratched across each well to create the scratch wound. Wells were aspirated to remove floating cells and replaced with 200 uL of culture media. The plate was placed in the Incucyte Live-Cell Imaging System (Essen BioScience), and images were collected hourly for 48 hours. Five wells of NS and 6 wells of each shATRX with clean scratch wounds were analyzed. Images were downloaded and analyzed by ImageJ (94). The MG-63 WT and KO cells were plated in the same way, however for these cells, we used the Woundmaker^TM^ (Essen BioScience) to create the scratch wounds in each well. Thirteen scratch-wound wells of WT and 10 wells of KO were analyzed in the same way as with the 143B cell line.

### Transwell migration and Boyden invasion assays

The Matrigel-coated invasion assay plate (Corning; catalog #354480) was removed from the foil bag and allowed to come to room temperature. Culture media consisting of DMEM and 1% Pen Strep, but no FBS, was also brought to room temperature, and then 500 µL of this media was added to each Matrigel-coated well to hydrate the inserts while incubating at 37°C for 4 hours. At this time, 1% BSA was added to 50 mLs of the DMEM culture media. The uncoated plate (Corning; catalog #354578) for the migration assay was removed from its foil bag. In each plate’s upper inserts, the 143B WT and KO cell lines were plated at 10,000 cells per well and the MG-63 WT and KO cell lines were plated at 20,000 cells per well in 200 µL of the DMEM with 1% BSA. Then 750 µL of DMEM with 1% Pen Strep and 10% FBS was placed in the bottom well below each insert. All plates were incubated at 37°C. The migration assay plates were removed after 18 hours and the Matrigel-coated plates were removed after 41 hours. Media was aspirated from the upper and lower chambers of each well and washed twice with 1x PBS. Cells were fixed with 4% paraformaldehyde for 5 minutes and then washed again with 1x PBS. Cells were then permeabilized with 750 µL Perm buffer (1x PBS with 0.2% Triton-X-100) and incubated on bench top for 20 minutes. Then the cells were washed again with 1X PBS and stained with 0.1% Crystal Violet for 20 minutes on bench top. Once again, cells were washed with 1x PBS and then non-migrated cells were scraped off the inside of each insert with a sterile cotton swab (VWR; catalog #149-0332). Inserts were allowed to dry for at least one hour. A scalpel was used to cut out each insert and mount to a microscope slide with Cytoseal (Thermo Scientific; catalog #83104) and a coverslip. Slides were dried for 3 hours and then imaged on a light microscope, recording five high powered fields per membrane. For the 143B cell line, 8 wells of WT and 8 wells of KO were analyzed for both the uncoated and Matrigel-coated assays. For the MG-63 cell line, 12 wells of WT and 12 wells of KO were analyzed for both uncoated and Matrigel-coated assays. Images were analyzed using ImageJ (94) to get total cell counts per slide.

### Osterix-Cre mouse model experiment

The conditional osteoblast-specific mouse model builds from a previously-established (36, 37) transgeneic *Osterix-Cre* (*Osx-Cre*) mouse model with conditional (floxed) alleles of both *p53* (*p53^fl/fl^*) and *Rb* (*Rb^fl/fl^*) and with a Tet-off cassette providing an additional level of temporal control. Female *Osx1-GFP::Cre* mice (mixed background) were kindly donated by the Karner lab, and male *Rb^fl/fl^p53^fl/fl^*mice (mixed background) as well as male *ATRX^fl/y^ Rb^fl/fl^p53^fl/fl^* mice (mixed background) were kindly donated by the Gibbons lab. These mice were crossed to generate experimental cohorts of males and females that were either *Osx-Cre+p53^fl/fl^Rb^fl/fl^* (n=26 males and 25 females) or *Osx-Cre+p53^fl/fl^Rb^fl/fl^ATRX^fl/fl/y^*(n=22 males and 25 females). Mice were fed a rodent diet supplemented with doxycycline until weaning (Envigo; catalog #TD.120769), upon which we removed the doxycycline diet and monitored mice of each genotype for tumor development. To more comprehensively identify tumors that developed in any bone and to further increase our sensitivity in tumor detection, we performed monthly fluoroscopy on a subcohort of 10 females and 10 males of each genotype (Supplemental Figure 1A). Tumors were collected, locations were noted, and gene recombination of tumors was confirmed by PCR to distinguish the knockout allele from the floxed (unrecombined) and wildtype allele (Supplemental Figures 1B).

### Subcutaneous xenografts for tumor growth comparison

For xenograft studies, SCID-beige mice (strain code 250) were obtained from Charles River Laboratories. For each cell line, we injected 1.0 x 10^6^ cells suspended in 200 μL saline subcutaneously into the flank of mice when they were 8 to 10 weeks old. For the shRNA/NS cells, there were three males and two females per treatment group, with three same-sex mice per cage and one mouse per treatment group (randomly chosen) in each cage. For the WT/KO cells, there were five males and five females per treatment group, with five same-sex mice per cage and two mice receiving one cell type and three mice receiving the other cell type (randomly chosen) per cage. Resultant tumors were measured every two to three days with calipers to compare growth rates. When the tumors grew to the point of humane endpoints (defined as respiratory changes, greater than fifteen percent body weight loss, dehydration, decreased physical activity, or large tumor burden), mice were sacrificed. Final tumor volume and mass were measured.

### Xenograft tumor immunohistochemistry

Xenograft tumors were harvested, formalin-fixed, and paraffin-embedded. The Duke Histopathology Core Research Group prepared slides and performed H&E and immunohistochemistry (IHC) staining for Ki-67, ATRX, and cleaved caspase 3. IHC staining was interpreted and quantified with the help of a board-certified veterinary pathologist.

### Orthotopic mouse model of metastasis-injections, amputations, IVIS imaging

The LUC-labelled 143B WT and CRISPR KO cell lines described below were prepared at a concentration of 6.0 x 10^5^ cells suspended in 30 μL 50:50 saline:Matrigel solution (Corning; catalog#354234). Cells were injected into the subperiosteal space of the tibia in 8- to 10-week-old SCID-beige mice (obtained from Duke University’s Division of Laboratory Animal Resources (DLAR) breeding core facility) (WT: n=5 males and 10 females, KO: n=6 males and 11 females). When tumors were 1 cm^3^ in size, the mice underwent amputation of the affected limb. Luciferase luminescence was measured with the IVIS imaging device immediately after injection and then every 2 to 3 days to compare metastasis burden and disease progression in our mice. For each imaging session, mice were anesthetized and then injected intraperitoneally with 100 µL of D-luciferin at 28.6 mg/mL (Gold Biotechnology; catalog #LUCNA-1G). IVIS images were collected sequentially for 30 minutes to capture peak luciferin tissue distribution. Luminescence readings were collected for the lung field region of interest for each mouse and compared to analyze metastatic burden in the lungs. The data shown in Figure 3K were gathered at one-week post-amputation.

### RNA Sequencing

RNA-Seq was performed on the *ATRX* knockdown or non-silenced 143B human OS cell lines in collaboration with the Duke Center for Genomic and Computational Biology. RNA-Seq data was processed using the TrimGalore toolkit which employs Cutadapt (95) to trim low quality bases and Illumina sequencing adapters from the 3’ end of the reads. Only reads that were 20nt or longer after trimming were kept for further analysis. Reads were mapped to the GRCh37v75 version of the human genome and transcriptome (96) using the STAR RNA-Seq alignment tool (97). Reads were kept for subsequent analysis if they mapped to a single genomic location. Gene counts were compiled using the HTSeq tool. Only genes that had at least ten reads in any given library were used in subsequent analysis. Normalization and differential expression were carried out using the DESeq2 Bioconductor package (98, 99) with the R statistical programming environment. The false discovery rate was calculated to control for multiple hypothesis testing. PCA results from sequencing samples are shown Supplemental Figure 5B. Gene set enrichment analysis (100) was performed to identify differentially regulated pathways and gene ontology terms for each of the comparisons performed.

### ATAC Sequencing

ATAC-Seq data was processed using the TrimGalore toolkit (http://www.bioinformatics.babraham.ac.uk/projects/trim_galore) which employs Cutadapt (95) to trim low quality bases and Illumina sequencing adapters from the 3’ end of the reads. Only reads that were 20nt or longer after trimming were kept for further analysis. Reads were mapped to the hg19 version of the human genome using the bowtie alignment tool (101). Reads were kept for subsequent analysis if they mapped to a single genomic location. Amplification artifacts were removed using the Picard toolkit (http://broadinstitute.github.io/picard). Regions of open chromatin regions were called using the MACS2 peak calling algorithm (102). The peaks were set to a 300nt window surrounding the mode location. Peaks that overlapped across the two conditions being compared were merged into a single peak. The number of reads from each individual sample that overlapped the peaks were quantified. Normalization and differential openness of chromatin regions were calculated using the DESeq2 Bioconductor package (98, 99) with the R statistical programming environment. Peaks were then annotated with their nearest gene using the GRCh37v758 of the human transcriptome. Quality control statistics are shown in Supplemental Figure 5C.

### Chromatin binding motif analysis

To understand ATRX chromatin binding, chromVAR was applied to analyze the ATAC-Seq data for variability of DNA motif enrichment in the accessible regions (103). In brief, chromVAR aggregates accessible regions sharing the same DNA motif, then compares the observed accessibility of all peaks containing that motif with a background set of peaks normalized for known technical confounders. The aligned fragments and peaks were used as inputs. For reading in peaks in the narrowpeak format, peaks were first resized to width = 500 bp and then in case of overlapping peaks, the peak with strongest signal was retained. In addition, peaks were filtered such that each peak should have at least one fragment across all the samples. To determine background peaks the GC content was used and hg19 genome sequence was selected as input. To identify significantly variable motifs among cells chromVAR scans the peaks for binding motif occurrences, using a curated collection of motifs provided from “motifmatchr” package (104). In addition to curated binding motifs, peaks were also scanned for all 6-mers. Cell-cell similarity was visualized in a two-dimensional t-SNE plot using the bias-corrected deviations in accessibility for both curated motifs and 6-mers.

### Nuclear extractions and NF-κB ELISA

143B and MG-63 wildtype and *ATRX* knockout cells were plated at 1 x 10^7^ cells per culture dish in two 10 cm culture dishes for each cell type. After 24 hours of incubation, cells were harvested and nuclear extractions were performed following the protocol of the Active Motif Nuclear Extract Kit (catalog #40010). Final extracted protein concentrations for each cell type were measured using a Bradford protein assay. The Active Motif TransAm NFκB Family kit (catalog #43296) was used to perform the ELISA assay with our nuclear extracts following the kit’s protocol. Recombinant p50 protein (Active Motif, catalog #31101) was used for the standard curve. Absorbance measurements were read using the SpectraMax i3x spectrophotometer.

### Bioactives compound screen

MG-63 wildtype and *ATRX* knockout cells were screened in triplicate with the Bioactive Compound Library (catalog #L1700) from Selleckchem, a diverse collection of 2100 compounds with demonstrated bioactivity. Library compounds were plated onto 384 well tissue culture plates using a Labcyte Echo for a final concentration of 1 uM after addition of 750 cells in 50 ul media using a Matrix WellMate. Cells were incubated with drug at 37°C for 72 hours and then assayed for viability using Cell Titer-Glo (Promega; catalog #G7571). Cell viability for each well was normalized against the DMSO wells contained in each plate. All screens were performed in the Duke Functional Genomics Shared Resource.

### IC^50^ assays

We further validated the integrin inhibitor SB273005 (Selleckchem; catalog #S7540) from the drug screen using IC^50^ assays. MG-63 wildtype and knockout cells were plated at 10,000 cells per well in 96-well plates with 100 uL DMEM. Cells were incubated overnight. Serial dilutions of drug or DMSO vehicle control were prepared to test plated concentrations of 1,000 nM, 200 nM, 40 nM, 8 nM, 1.6 nM and 0.32 nM drug. After incubation for 24 hours, 80 uL Cell Titer-Glo reagent (Promega, catalog #G7571) was added to each well using a multichannel pipette. The plate was wrapped in aluminum foil and placed on an orbital shaker for five minutes. Then, the plate was imaged on the SpectraMax i3x spectrophotometer at ten minutes, thirty minutes, and one hour.

### *In vivo* study with integrin inhibitor SB273005

The integrin-inhibitor SB273005 (Selleckchem; catalog #S7540) was tested *in vivo* with xenograft mouse tumors to further validate its efficacy as a therapeutic for *ATRX*-deficient OS. We used the U-2OS human OS cell line (ATRX-null) to create subcutaneous tumors in NSG mice (obtained from Duke University’s DLAR rodent breeding core facility), injecting 1.0 x 10^6^ cells in 200 μL of a 50:50 saline:Matrigel solution subcutaneously in the flank. Each cohort contained five male and five female mice at 8 to 10 weeks of age. As tumors became palpable, mice were randomly assigned to treatment with a vehicle control or SB273005 at a dose of 50 mg/kg/day, a dose well-tolerated and therapeutically effective in rats in a study by Badger et al. (43). Tumor volume was monitored with caliper measurements every two to three days for a thirty-day treatment course.

### *In vitro* assays with integrin inhibitor SB273005

Cell Titer-Glo fluorescence assays of cell viability were repeated, and a dose of 8 nM was determined to be the highest dose of SB273005 (Selleckchem; catalog #S7540) that could be administered to cells *in vitro* with minimal differential cell survival between wildtype and knockout MG-63 cells. Cells were plated and treated with SB273005 or DMSO (vehicle) alone at the 8 nM dose, and *in vitro* scratch wound assays and migration and invasion chamber assays were repeated as previously described, but with four treatment groups: wildtype cells treated with vehicle control, wildtype cells treated with integrin inhibitor, knockout cells treated with vehicle control, and knockout cells treated with integrin inhibitor.

### Analysis of the ICGC/TCGA Pan-Cancer Analysis of Whole Genomes sequencing dataset

cBioPortal was used to compare *ATRX*-altered versus *ATRX*-unaltered tumors in the ICGC/TCGA Pan-Cancer Analysis of Whole Genomes dataset (18, 19, 44). An oncoPrint analysis was perfomed on the cBioPortal query while filtering for mutations and copy number alterations of known significance. In the (n=21) altered group, two patients have missense mutations, one patient has an amplification, and the remaining patients have *ATRX* deletions. Over-representation Analysis (ORA) was then performed on expression data between *ATRX*-altered and unaltered groups (105). Genes submitted to ORA had higher expression in *ATRX*-altered groups than unaltered groups, and all identified genes had q values < 0.05. The query was submitted to the Panther, KEGG, and Wikipathway cancer databases.

### Quantification and Statistical Analysis

For bar graphs, all data are presented as means +/-SEM. All data were analyzed for statistically significant differences using the Student’s t-test (for two comparisons) or analysis of variance with Tukey’s multiple comparisons test (for multiple comparisons). Effects over time were analyzed with repeated-measures analysis of variance. Tumor-free survival curves were estimated using the Kaplan-Meier method and compared statistically using the log-rank test. Next generation sequencing statistical analysis is described in the RNA-Seq and ATAC-Seq methods section. JMP Pro 15.0 and/or Prism 9.0 (GraphPad Software, Inc.) were used for the statistical analyses. Any p value less than 0.05 was considered statistically significant.

### Study Approval

All animal studies were approved by the Institutional Animal Care and Use Committee (IACUC) at Duke University.

### Materials & Correspondence

Further information and requests for resources and reagents should be directed to and will be fulfilled by the Lead Contact, William C. Eward.

### Data Availability

The high-throughput sequencing data from this study have been submitted to the NCBI GEO repository under accession number GSE167546.

### Author Contributions

SBD, WCE, BA, and JAS conceived the project, designed experiments, and wrote the manuscript. SBD analyzed data. SBD and SHP bred, managed, and genotyped mice. Immunofluorescence was performed by HB. SBD and DS performed experiments with the integrin inhibitor SB273005 both *in vitro* and *in vivo*. SBD, EJM, and WF performed genotyping of subcutaneous tumors to validate *ATRX* recombination. EJM also assisted with cross-referencing gene expression data and chromatin accessibility changes in the RNA-Seq and ATAC-Seq data. MZ and JR assisted with the common binding motif analysis. VK and SV studied growth of cells on Matrigel beds. JS and NH analyzed sequencing data from cBioPortal. WF and DGK provided consultation on experimental design. MUS provided the graphic of tumor phenotypes of aggression.

## Acknowledgements

We thank Richard Gibbons for providing the *ATRX^fl/fl^* mice and Anton Berns for providing the *Rb^fl/fl^p53^fl/fl^* mice. We thank Courtney Karner for providing the *Osterix-Cre* mice. This work was supported by the Consortium for Canine Comparative Oncology (C3O), Hyundai Hope on Wheels, and the Duke Health Scholars Award.

---

**Supplemental Figure 1:**
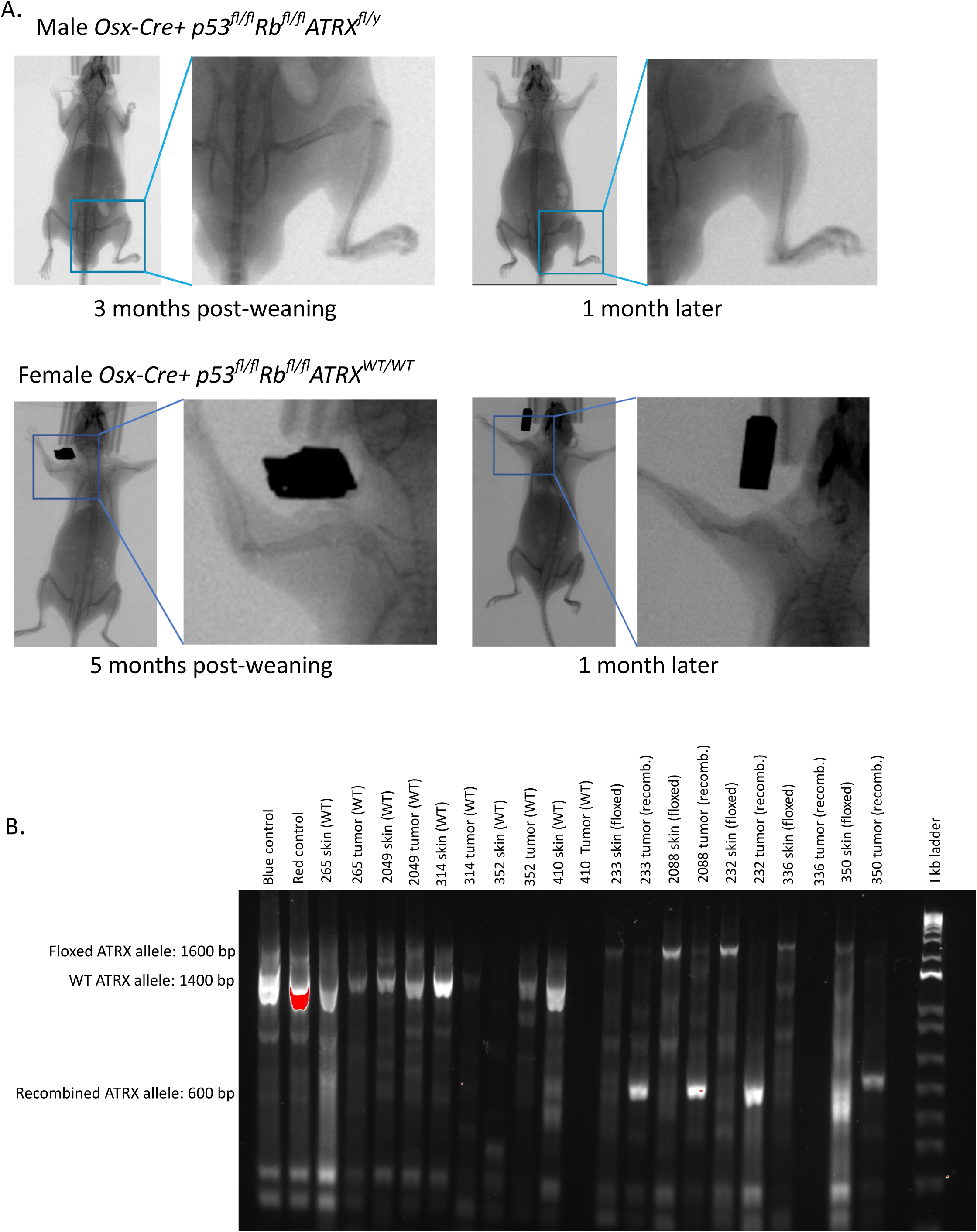
Loss of *ATRX* expression increased the rate of tumor formation in an *Osterix-Cre* driven mouse model of OS. A) Examples of fluoroscopic imaging to detect tumors in the Osx-Cre driven mouse model. Top: This male *ATRX*-floxed mouse developed an OS of the distal femur at 3 months post-weaning. This tumor was not palpable on physical examination at this stage but was detectable by fluoroscopy. One month later, this tumor had grown and was now detectable on physical examination as well. Bottom: This female *ATRX*-WT mouse developed an OS of the proximal humerus at 5 months post-weaning. Again, this tumor was detected by fluoroscopy, but was first palpable on physical examination one month later. B) Genotyping was used to validate *ATRX* recombination or wildtype status with DNA collected from a subset of mouse tumors with matched skin samples. Primers used were PPS 1.17 5’-AAC TCA TTC AAC TGC CC-3’ and PPS 1.28 5’-CAT TTA ATC CCT CCT GCC-3’. WT band at 1400 bp, ATRX floxed band at 1600 bp, and ATRX-NEO recombined band at 600 bp.

**Supplemental Figure 2:**
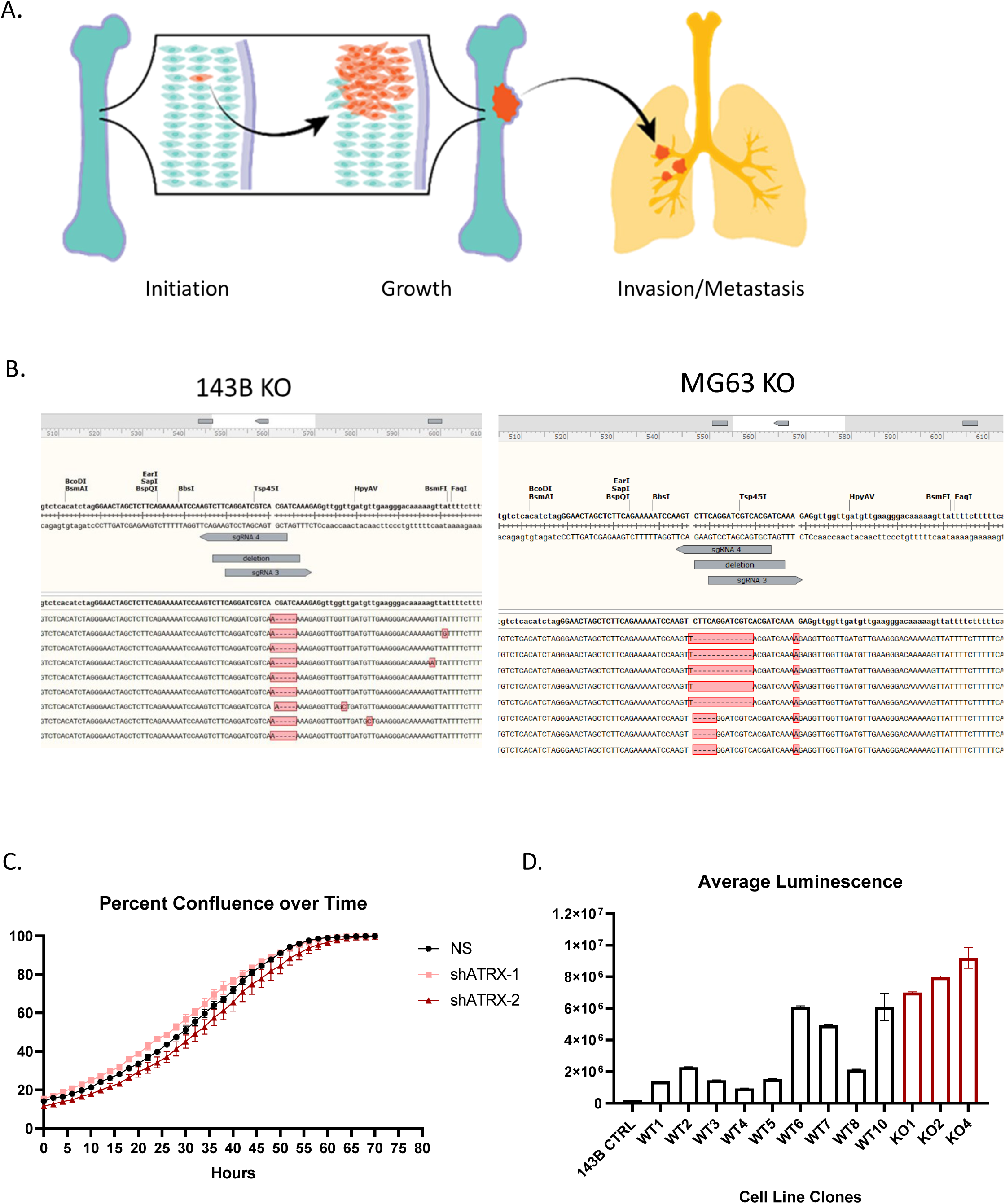
A) Aggressive tumor cellular phenotypes, including tumor initiation, tumor growth, and migration, invasion and metastasis. B) Sanger sequencing results showing effective CRISPR-Cas9 knockout of *ATRX* expression in the 143B human OS cell line and the MG-63 human OS cell line. C) All 143B cell types (NS and shRNA KDs) showed high cellular proliferation with no significant difference in measure of percent confluence over time detected with the Incucyte Live Cell Imager. D) Comparison of average luminescence in individual 143B WT and KO clones transfected with the LUC plasmid. WT clone 6 and KO clone 1 were similar in luminescence levels, so these were used for the orthotopic injection experiment.

**Supplemental Figure 3:**
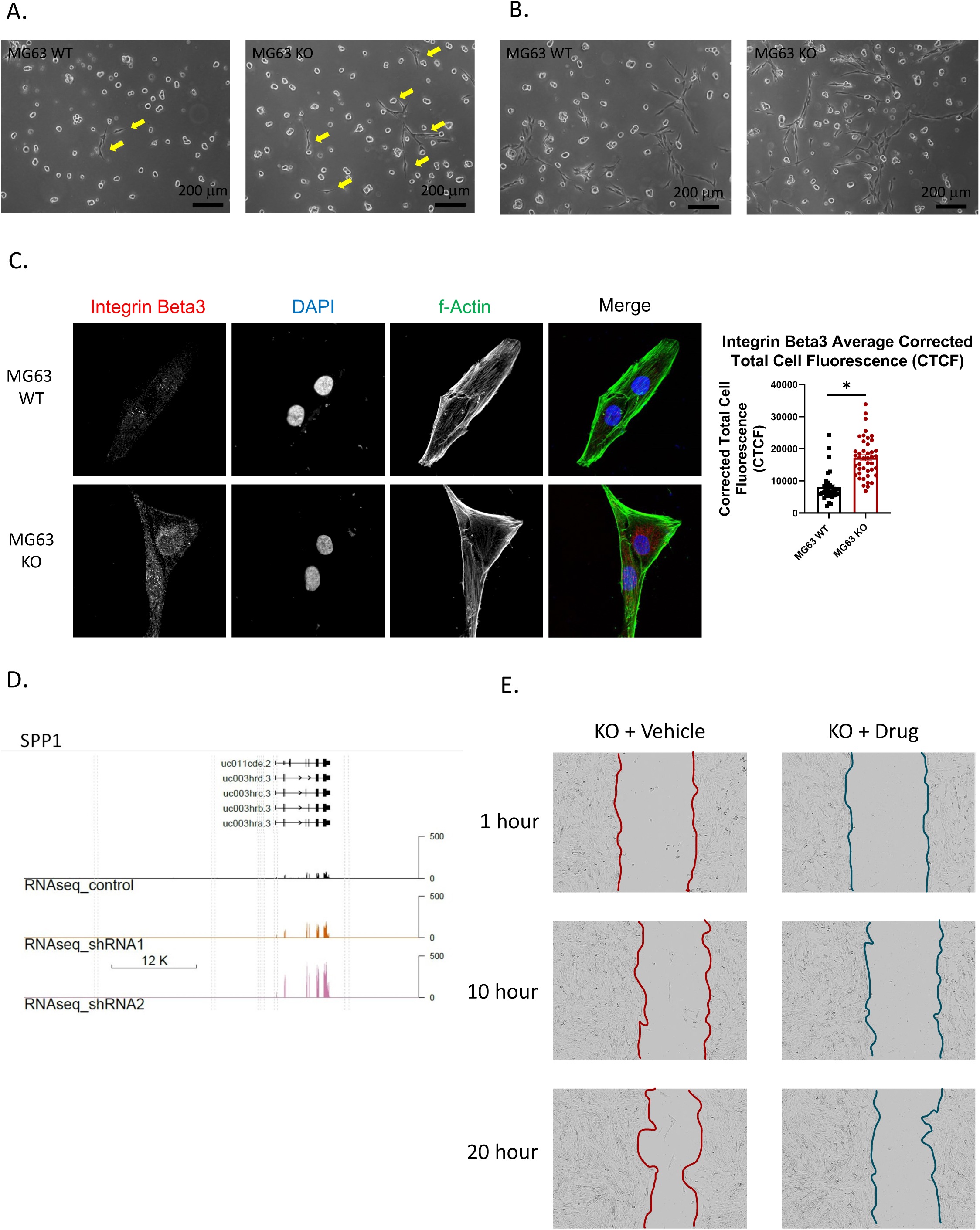
A) and B) Notable differences in morphological appearances (specifically, increased branching networks between cells) were apparent between WT and KO cells in the slower-growing MG-63 cell line after 72 hours (A) and 96 hours (B) Yellow arrows highlight the increased branching networks forming between KO cells relative to WT cells at 72 hours C) MG-63 *ATRX* KO cells show significantly increased expression of integrin β3 compared to WT cells (Student’s t-test, p<0.0001, KO n=43, WT n=35). D) RNA-Seq gene track for osteopontin (*SPP1*) showing increased mRNA expression with ATRX KD (log fold change over NS control of 1.3 and 2.52 for shATRX-1 and shATRX-2, respectively). Osteopontin is one of the key matrix components to which integrins αvβ3 bind, and expression of osteopontin has been correlated with poor survival in patients with OS. E) Representative images of KO+Vehicle compared to KO+Drug scratch wounds at three time points in the MG-63 cell line treated with 8nM SB273005 or vehicle control (DMSO).

**Supplemental Figure 4:**
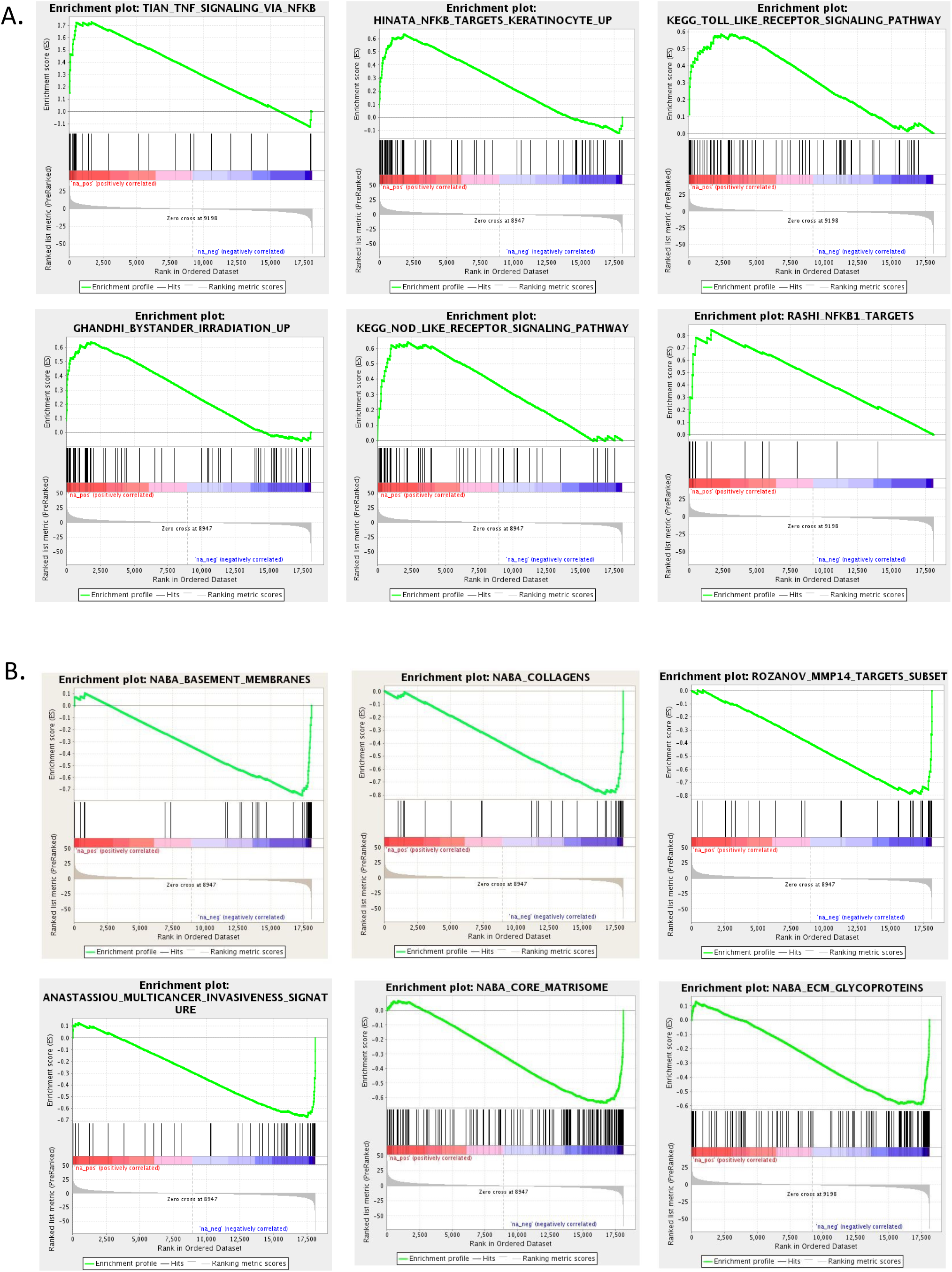
Enrichment plots for GSEA pathways altered with *ATRX* KD. A) Enrichment plots for several of the most upregulated pathways found by GSEA (Curated Pathways). Many of these pathways were associated with NF-κB signaling. B) Enrichment plots for several of the most downregulated pathways found by GSEA (Curated Pathways). Many of these pathways are related to ECM remodeling.

**Supplemental Figure 5:**
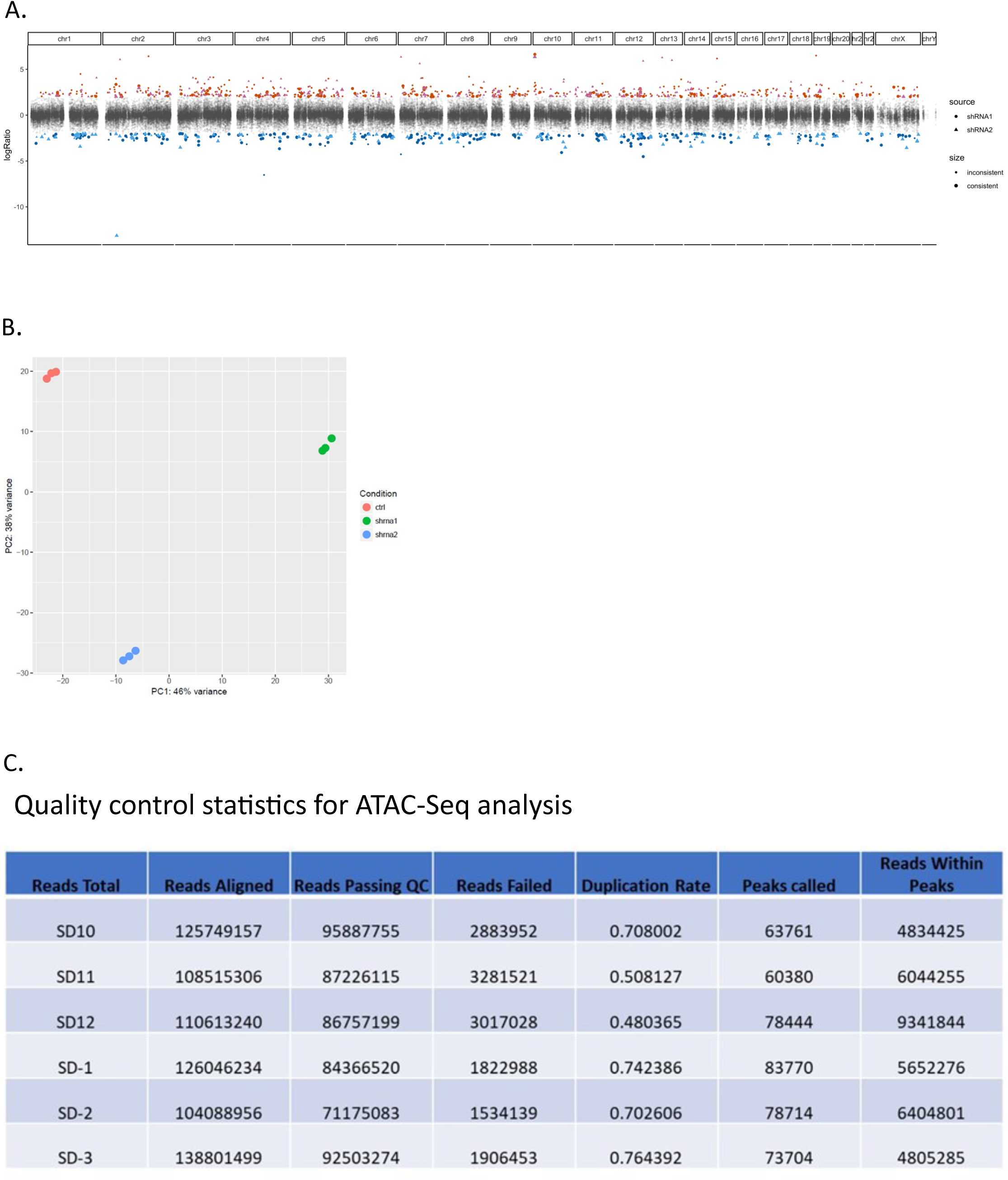
A) Manhattan plot of ATAC-Seq regions of differential chromatin openness for each shRNA compared to non-silenced control. These data support a global genomic role of *ATRX* as a chromatin remodeler. B) PCA results as quality check of samples used for RNA sequencing. C) Quality control statistics for ATAC-Seq analysis.

**Supplemental Figure 6:**
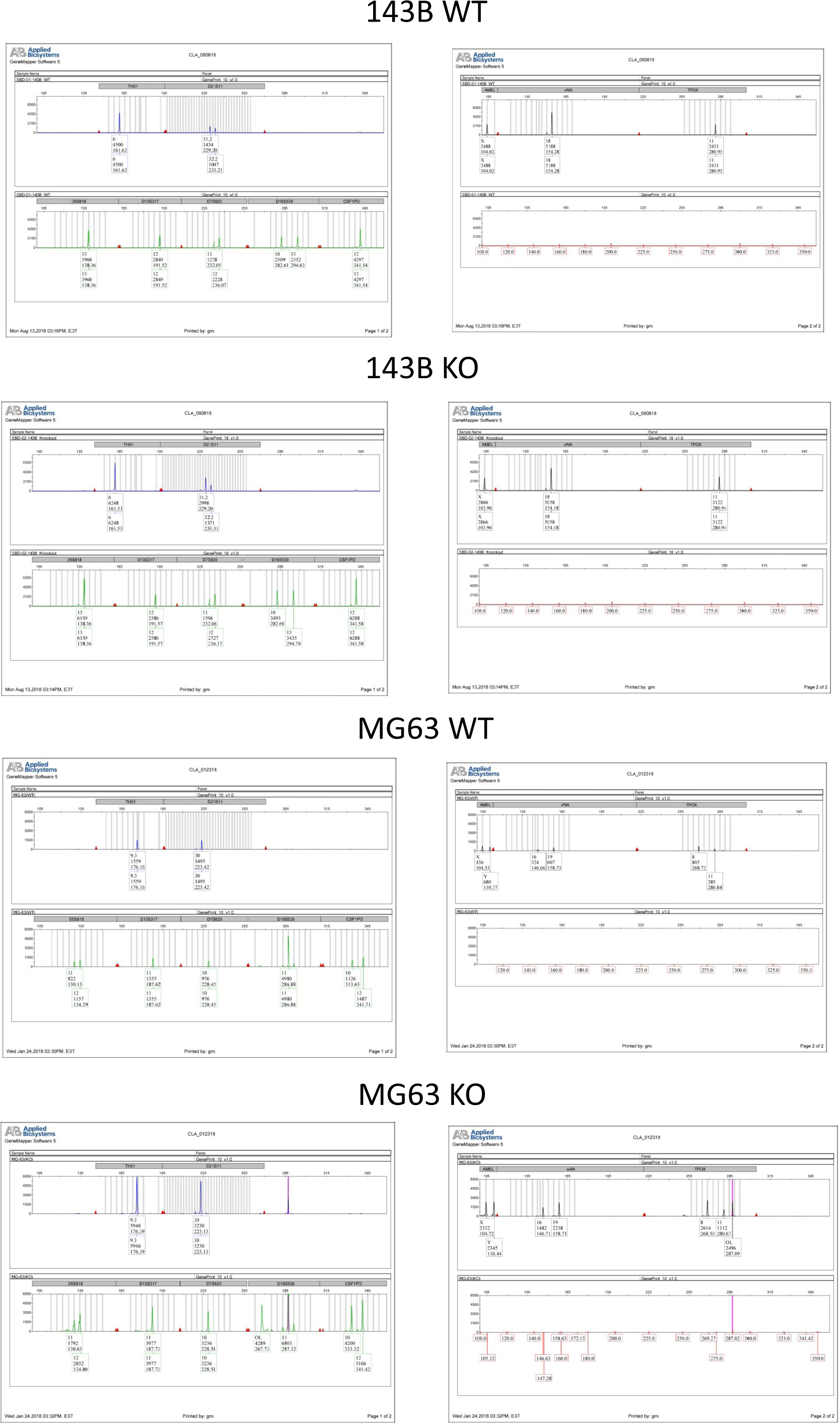
Chromatographs verifying cell line authenticity for all cell lines used in this manuscript.

## References

1. Ottaviani G, Jaffe N. The epidemiology of osteosarcoma. Cancer Treat Res. 2009;152:3–13. doi:10.1007/978-1-4419-0284-9_1

2. Mirabello L, Troisi RJ, Savage SA. International osteosarcoma incidence patterns in children and adolescents, middle ages and elderly persons. Int J Cancer. Jul 1 2009;125(1):229–34. doi:10.1002/ijc.24320

3. Bielack SS, et al. Prognostic factors in high-grade osteosarcoma of the extremities or trunk: an analysis of 1,702 patients treated on neoadjuvant cooperative osteosarcoma study group protocols. J Clin Oncol. Feb 1 2002;20(3):776–90. doi:10.1200/JCO.2002.20.3.776

4. Stephens PJ, et al. Massive genomic rearrangement acquired in a single catastrophic event during cancer development. Cell. Jan 7 2011;144(1):27–40. doi:10.1016/j.cell.2010.11.055

5. Chen X, et al. Recurrent somatic structural variations contribute to tumorigenesis in pediatric osteosarcoma. Cell Rep. Apr 10 2014;7(1):104–12. doi:10.1016/j.celrep.2014.03.003

6. Bousquet M, et al. Whole-exome sequencing in osteosarcoma reveals important heterogeneity of genetic alterations. Ann Oncol. Apr 2016;27(4):738–44. doi:10.1093/annonc/mdw009

7. Rickel K, Fang F, Tao J. Molecular genetics of osteosarcoma. Bone. Sep 2017;102:69–79. doi:10.1016/j.bone.2016.10.017

8. Perry JA, et al. Complementary genomic approaches highlight the PI3K/mTOR pathway as a common vulnerability in osteosarcoma. Proc Natl Acad Sci U S A. Dec 23 2014;111(51):E5564–73. doi:10.1073/pnas.1419260111

9. Kovac M, et al. Exome sequencing of osteosarcoma reveals mutation signatures reminiscent of BRCA deficiency. Nat Commun. Dec 3 2015;6:8940. doi:10.1038/ncomms9940

10. Jiao Y, et al. Frequent ATRX, CIC, FUBP1 and IDH1 mutations refine the classification of malignant gliomas. Oncotarget. Jul 2012;3(7):709–22. doi:10.18632/oncotarget.588

11. Koelsche C, et al. Differential nuclear ATRX expression in sarcomas. Histopathology. Apr 2016;68(5):738–45. doi:10.1111/his.12812

12. Qadeer ZA, et al. Decreased expression of the chromatin remodeler ATRX associates with melanoma progression. J Invest Dermatol. Jun 2014;134(6):1768–1772. doi:10.1038/jid.2014.45

13. Jiao Y, et al. DAXX/ATRX, MEN1, and mTOR pathway genes are frequently altered in pancreatic neuroendocrine tumors. Science. Mar 4 2011;331(6021):1199–203. doi:10.1126/science.1200609

14. Liau JY, et al. Leiomyosarcoma with alternative lengthening of telomeres is associated with aggressive histologic features, loss of ATRX expression, and poor clinical outcome. Am J Surg Pathol. Feb 2015;39(2):236–44. doi:10.1097/PAS.0000000000000324

15. Huether R, et al. The landscape of somatic mutations in epigenetic regulators across 1,000 paediatric cancer genomes. Nat Commun. Apr 8 2014;5:3630. doi:10.1038/ncomms4630

16. Danussi C, et al. Atrx inactivation drives disease-defining phenotypes in glioma cells of origin through global epigenomic remodeling. Nat Commun. Mar 13 2018;9(1):1057. doi:10.1038/s41467-018-03476-6

17. Cancer Genome Atlas Research N, et al. The Cancer Genome Atlas Pan-Cancer analysis project. Nat Genet. Oct 2013;45(10):1113–20. doi:10.1038/ng.2764

18. Cerami E, et al. The cBio cancer genomics portal: an open platform for exploring multidimensional cancer genomics data. Cancer Discov. May 2012;2(5):401–4. doi:10.1158/2159-8290.CD-12-0095

19. Gao J, et al. Integrative analysis of complex cancer genomics and clinical profiles using the cBioPortal. Sci Signal. Apr 2 2013;6(269):pl1. doi:10.1126/scisignal.2004088

20. Consortium APG. AACR Project GENIE: Powering Precision Medicine through an International Consortium. Cancer Discov. Aug 2017;7(8):818–831. doi:10.1158/2159-8290.CD-17-0151

21. Liau JY, et al. Comprehensive screening of alternative lengthening of telomeres phenotype and loss of ATRX expression in sarcomas. Mod Pathol. Dec 2015;28(12):1545–54. doi:10.1038/modpathol.2015.114

22. Kreilmeier T, et al. Alternative lengthening of telomeres does exist in various canine sarcomas. Mol Carcinog. Mar 2017;56(3):923–935. doi:10.1002/mc.22546

23. Ye K, et al. Systematic discovery of complex insertions and deletions in human cancers. Nat Med. Jan 2016;22(1):97–104. doi:10.1038/nm.4002

24. Gibbons R. Alpha thalassaemia-mental retardation, X linked. Orphanet J Rare Dis. May 4 2006;1:15. doi:10.1186/1750-1172-1-15

25. Argentaro A, et al. Structural consequences of disease-causing mutations in the ATRX-DNMT3-DNMT3L (ADD) domain of the chromatin-associated protein ATRX. Proc Natl Acad Sci U S A. Jul 17 2007;104(29):11939–44. doi:10.1073/pnas.0704057104

26. Clynes D, Higgs DR, Gibbons RJ. The chromatin remodeller ATRX: a repeat offender in human disease. Trends Biochem Sci. Sep 2013;38(9):461–6. doi:10.1016/j.tibs.2013.06.011

27. Voon HP, Wong LH. New players in heterochromatin silencing: histone variant H3.3 and the ATRX/DAXX chaperone. Nucleic Acids Res. Feb 29 2016;44(4):1496–501. doi:10.1093/nar/gkw012

28. Shi L, Wen H, Shi X. The Histone Variant H3.3 in Transcriptional Regulation and Human Disease. J Mol Biol. Jun 30 2017;429(13):1934–1945. doi:10.1016/j.jmb.2016.11.019

29. Napier CE, et al. ATRX represses alternative lengthening of telomeres. Oncotarget. Jun 30 2015;6(18):16543–58. doi:10.18632/oncotarget.3846

30. Lovejoy CA, et al. Loss of ATRX, genome instability, and an altered DNA damage response are hallmarks of the alternative lengthening of telomeres pathway. PLoS Genet. 2012;8(7):e1002772. doi:10.1371/journal.pgen.1002772

31. Hanahan D, Weinberg RA. Hallmarks of cancer: the next generation. Cell. Mar 4 2011;144(5):646–74. doi:10.1016/j.cell.2011.02.013

32. Haase S, et al. Mutant ATRX: uncovering a new therapeutic target for glioma. Expert Opin Ther Targets. Jul 2018;22(7):599–613. doi:10.1080/14728222.2018.1487953

33. Liang J, et al. Global changes in chromatin accessibility and transcription following ATRX inactivation in human cancer cells. FEBS Lett. Jan 2020;594(1):67–78. doi:10.1002/1873-3468.13549

34. Trojanowska M. Ets factors and regulation of the extracellular matrix. Oncogene. Dec 18 2000;19(55):6464–71. doi:10.1038/sj.onc.1204043

35. Oikawa T, Yamada T. Molecular biology of the Ets family of transcription factors. Gene. Jan 16 2003;303:11–34. doi:10.1016/s0378-1119(02)01156-3

36. Berman SD, et al. Metastatic osteosarcoma induced by inactivation of Rb and p53 in the osteoblast lineage. Proc Natl Acad Sci U S A. Aug 19 2008;105(33):11851–6. doi:10.1073/pnas.0805462105

37. Walkley CR, et al. Conditional mouse osteosarcoma, dependent on p53 loss and potentiated by loss of Rb, mimics the human disease. Genes Dev. Jun 15 2008;22(12):1662–76. doi:10.1101/gad.1656808

38. Rodda SJ, McMahon AP. Distinct roles for Hedgehog and canonical Wnt signaling in specification, differentiation and maintenance of osteoblast progenitors. Development. Aug 2006;133(16):3231–44. doi:10.1242/dev.02480

39. Ricci B, et al. Osterix-Cre marks distinct subsets of CD45- and CD45+ stromal populations in extra-skeletal tumors with pro-tumorigenic characteristics. Elife. Aug 5 2020;9doi:10.7554/eLife.54659

40. Chen J, et al. Osx-Cre targets multiple cell types besides osteoblast lineage in postnatal mice. PLoS One. 2014;9(1):e85161. doi:10.1371/journal.pone.0085161

41. Reddy GB, et al. Preclinical Testing of a Novel Niclosamide Stearate Prodrug Therapeutic (NSPT) Shows Efficacy Against Osteosarcoma. Mol Cancer Ther. Jul 2020;19(7):1448–1461. doi:10.1158/1535-7163.MCT-19-0689

42. Miller WH, et al. Discovery of orally active nonpeptide vitronectin receptor antagonists based on a 2-benzazepine Gly-Asp mimetic. J Med Chem. Jan 13 2000;43(1):22–6. doi:10.1021/jm990446u

43. Badger AM, et al. Disease-modifying activity of SB 273005, an orally active, nonpeptide alphavbeta3 (vitronectin receptor) antagonist, in rat adjuvant-induced arthritis. Arthritis Rheum. Jan 2001;44(1):128–37. doi:10.1002/1529-0131(200101)44:1<128::AID-ANR17>3.0.CO;2-M

44. Consortium ITP-CAoWG. Pan-cancer analysis of whole genomes. Nature. Feb 2020;578(7793):82-93. doi:10.1038/s41586-020-1969-6

45. Karin M. Nuclear factor-kappaB in cancer development and progression. Nature. May 25 2006;441(7092):431-6. doi:10.1038/nature04870

46. Xia Y, Shen S, Verma IM. NF-kappaB, an active player in human cancers. Cancer Immunol Res. Sep 2014;2(9):823–30. doi:10.1158/2326-6066.CIR-14-0112

47. Huber MA, et al. NF-kappaB is essential for epithelial-mesenchymal transition and metastasis in a model of breast cancer progression. J Clin Invest. Aug 2004;114(4):569–81. doi:10.1172/JCI21358

48. Felx M, et al. Endothelin-1 (ET-1) promotes MMP-2 and MMP-9 induction involving the transcription factor NF-kappaB in human osteosarcoma. Clin Sci (Lond). Jun 2006;110(6):645–54. doi:10.1042/CS20050286

49. Zhao Z, et al. Downregulation of MCT1 inhibits tumor growth, metastasis and enhances chemotherapeutic efficacy in osteosarcoma through regulation of the NF-kappaB pathway. Cancer Lett. Jan 1 2014;342(1):150–8. doi:10.1016/j.canlet.2013.08.042

50. Scatena M, et al. NF-kappaB mediates alphavbeta3 integrin-induced endothelial cell survival. J Cell Biol. May 18 1998;141(4):1083–93. doi:10.1083/jcb.141.4.1083

51. Broadhead ML, et al. The molecular pathogenesis of osteosarcoma: a review. Sarcoma. 2011;2011:959248. doi:10.1155/2011/959248

52. Kwakwa KA, Sterling JA. Integrin alphavbeta3 Signaling in Tumor-Induced Bone Disease. Cancers (Basel). Jul 8 2017;9(7)doi:10.3390/cancers9070084

53. Li R, et al. NF-kappaB signaling and integrin-beta1 inhibition attenuates osteosarcoma metastasis via increased cell apoptosis. Int J Biol Macromol. Feb 15 2019;123:1035–1043. doi:10.1016/j.ijbiomac.2018.11.003

54. Shi K, et al. Clinicopathological and prognostic values of fibronectin and integrin alphavbeta3 expression in primary osteosarcoma. World J Surg Oncol. Jan 28 2019;17(1):23. doi:10.1186/s12957-019-1566-z

55. Wang S, et al. SB-273005, an antagonist of alphavbeta3 integrin, reduces the production of Th2 cells and cytokine IL-10 in pregnant mice. Exp Ther Med. Jun 2014;7(6):1677–1682. doi:10.3892/etm.2014.1667

56. Gomes N, et al. Breast adenocarcinoma cell adhesion to the vascular subendothelium in whole blood and under flow conditions: effects of alphavbeta3 and alphaIIbbeta3 antagonists. Clin Exp Metastasis. 2004;21(6):553–61. doi:10.1007/s10585-004-3756-4

57. Reardon DA, et al. Randomized phase II study of cilengitide, an integrin-targeting arginine-glycine-aspartic acid peptide, in recurrent glioblastoma multiforme. J Clin Oncol. Dec 1 2008;26(34):5610–7. doi:10.1200/JCO.2008.16.7510

58. Stupp R, et al. Cilengitide combined with standard treatment for patients with newly diagnosed glioblastoma with methylated MGMT promoter (CENTRIC EORTC 26071-22072 study): a multicentre, randomised, open-label, phase 3 trial. Lancet Oncol. Sep 2014;15(10):1100–8. doi:10.1016/S1470-2045(14)70379-

59. Haddad T, et al. A phase I study of cilengitide and paclitaxel in patients with advanced solid tumors. Cancer Chemother Pharmacol. Jun 2017;79(6):1221–1227. doi:10.1007/s00280-017-3322-9

60. Landen CN, et al. Tumor-selective response to antibody-mediated targeting of alphavbeta3 integrin in ovarian cancer. Neoplasia. Nov 2008;10(11):1259–67. doi:10.1593/neo.08740

61. Hersey P, et al. A randomized phase 2 study of etaracizumab, a monoclonal antibody against integrin alpha(v)beta(3), + or -dacarbazine in patients with stage IV metastatic melanoma. Cancer. Mar 15 2010;116(6):1526–34. doi:10.1002/cncr.24821

62. Cirkel GA, et al. A dose escalating phase I study of GLPG0187, a broad spectrum integrin receptor antagonist, in adult patients with progressive high-grade glioma and other advanced solid malignancies. Invest New Drugs. Apr 2016;34(2):184–92. doi:10.1007/s10637-015-0320-9

63. Arun B, et al. The PARP inhibitor AZD2281 (Olaparib) induces autophagy/mitophagy in BRCA1 and BRCA2 mutant breast cancer cells. Int J Oncol. Jul 2015;47(1):262–8. doi:10.3892/ijo.2015.3003

64. Boldrini L, et al. Prognostic significance of osteopontin expression in early-stage non-small-cell lung cancer. Br J Cancer. Aug 22 2005;93(4):453–7. doi:10.1038/sj.bjc.6602715

65. El-Tanani MK, et al. The regulation and role of osteopontin in malignant transformation and cancer. Cytokine Growth Factor Rev. Dec 2006;17(6):463–74. doi:10.1016/j.cytogfr.2006.09.010

66. Forootan SS, et al. Prognostic significance of osteopontin expression in human prostate cancer. Int J Cancer. May 1 2006;118(9):2255–61. doi:10.1002/ijc.21619

67. Song JY, et al. Osteopontin expression correlates with invasiveness in cervical cancer. Aust N Z J Obstet Gynaecol. Aug 2009;49(4):434–8. doi:10.1111/j.1479-828X.2009.01027.x

68. Shevde LA, Samant RS. Role of osteopontin in the pathophysiology of cancer. Matrix Biol. Jul 2014;37:131–41. doi:10.1016/j.matbio.2014.03.001

69. Anborgh PH, et al. Role of plasma osteopontin as a biomarker in locally advanced breast cancer. Am J Transl Res. 2015;7(4):723–32.

70. Gu X, et al. Prognostic significance of osteopontin expression in gastric cancer: a meta-analysis. Oncotarget. Oct 25 2016;7(43):69666–69673. doi:10.18632/oncotarget.11936

71. Liu F, Bai C, Guo Z. The prognostic value of osteopontin in limited-stage small cell lung cancer patients and its mechanism. Oncotarget. Sep 19 2017;8(41):70084–70096. doi:10.18632/oncotarget.19589

72. Wang H, et al. Increased expression of osteopontin indicates poor prognosis in hepatocellular carcinoma. Int J Clin Exp Pathol. 2018;11(12):5916–5922.

73. Pang X, et al. Osteopontin as a multifaceted driver of bone metastasis and drug resistance. Pharmacol Res. Jun 2019;144:235–244. doi:10.1016/j.phrs.2019.04.030

74. Gaumann A, et al. Osteopontin expression in primary sarcomas of the pulmonary artery. Virchows Arch. Nov 2001;439(5):668–74. doi:10.1007/s004280100452

75. Song K, et al. Regulation of osteosarcoma cell invasion through osteopontin modification by miR-4262. Tumour Biol. May 2016;37(5):6493–9. doi:10.1007/s13277-015-4530-8

76. Pal M. Tumor metastasis suppressor functions of Ets transcription factor through integrin beta3-mediated signaling pathway. J Cell Physiol. Nov 2019;234(11):20266–20274. doi:10.1002/jcp.28627

77. Li R, Pei H, Watson DK. Regulation of Ets function by protein - protein interactions. Oncogene. Dec 18 2000;19(55):6514–23. doi:10.1038/sj.onc.1204035

78. Rothhammer T, et al. The Ets-1 transcription factor is involved in the development and invasion of malignant melanoma. Cell Mol Life Sci. Jan 2004;61(1):118–28. doi:10.1007/s00018-003-3337-8

79. Hahne JC, et al. Ets-1 expression promotes epithelial cell transformation by inducing migration, invasion and anchorage-independent growth. Oncogene. Aug 11 2005;24(34):5384–8. doi:10.1038/sj.onc.1208761

80. Hoesel B, Schmid JA. The complexity of NF-kappaB signaling in inflammation and cancer. Mol Cancer. Aug 2 2013;12:86. doi:10.1186/1476-4598-12-86

81. Longoni N, et al. ETS transcription factor ESE1/ELF3 orchestrates a positive feedback loop that constitutively activates NF-kappaB and drives prostate cancer progression. Cancer Res. Jul 15 2013;73(14):4533–47. doi:10.1158/0008-5472.CAN-12-4537

82. Dittmer J. The role of the transcription factor Ets1 in carcinoma. Semin Cancer Biol. Dec 2015;35:20–38. doi:10.1016/j.semcancer.2015.09.010

83. Wernert N, et al. Stromal expression of c-Ets1 transcription factor correlates with tumor invasion. Cancer Res. Nov 1 1994;54(21):5683–8.

84. Donahue JP, Sugg N, Hawiger J. The integrin alpha v gene: identification and characterization of the promoter region. Biochim Biophys Acta. Sep 13 1994;1219(1):228–32. doi:10.1016/0167-4781(94)90278-x

85. Sato M, et al. Transcriptional regulation of osteopontin gene in vivo by PEBP2alphaA/CBFA1 and ETS1 in the skeletal tissues. Oncogene. Sep 24 1998;17(12):1517–25. doi:10.1038/sj.onc.1202064

86. Oda N, Abe M, Sato Y. ETS-1 converts endothelial cells to the angiogenic phenotype by inducing the expression of matrix metalloproteinases and integrin beta3. J Cell Physiol. Feb 1999;178(2):121–32. doi:10.1002/(SICI)1097-4652(199902)178:2<121::AID-JCP1>3.0.CO;2-F

87. Tajima A, et al. Mouse integrin alphav promoter is regulated by transcriptional factors Ets and Sp1 in melanoma cells. Biochim Biophys Acta. Jul 24 2000;1492(2-3):377–84. doi:10.1016/s0167-4781(00)00121-4

88. Fang LW, et al. Ets-1 enhances tumor migration through regulation of CCR7 expression. BMB Rep. Sep 2019;52(9):548–553.

89. Li R, et al. EAP1/Daxx interacts with ETS1 and represses transcriptional activation of ETS1 target genes. Oncogene. Feb 10 2000;19(6):745–53. doi:10.1038/sj.onc.1203385

90. Xue Y, et al. The ATRX syndrome protein forms a chromatin-remodeling complex with Daxx and localizes in promyelocytic leukemia nuclear bodies. Proc Natl Acad Sci U S A. Sep 16 2003;100(19):10635–40. doi:10.1073/pnas.1937626100

91. Dyer MA, et al. ATRX and DAXX: Mechanisms and Mutations. Cold Spring Harb Perspect Med. Mar 1 2017;7(3)doi:10.1101/cshperspect.a026567

92. Montague TG, et al. CHOPCHOP: a CRISPR/Cas9 and TALEN web tool for genome editing. Nucleic Acids Res. Jul 2014;42(Web Server issue):W401-7. doi:10.1093/nar/gku410

93. Bae S, Park J, Kim JS. Cas-OFFinder: a fast and versatile algorithm that searches for potential off-target sites of Cas9 RNA-guided endonucleases. Bioinformatics. May 15 2014;30(10):1473–5. doi:10.1093/bioinformatics/btu048

94. ImageJ. U. S. National Institutes of Health, Bethesda, MD, USA; 1997-2018. https://imagej.nih.gov/ij/

95. Martin M. CutAdapt removes adapter sequences from high-throughput sequencing reads. Bioinformatics in action. 2011;17:10–12.

96. Kersey PJ, et al. Ensembl Genomes: an integrative resource for genome-scale data from non-vertebrate species. Nucleic Acids Res. Jan 2012;40(Database issue):D91-7. doi:10.1093/nar/gkr895

97. Dobin A, et al. STAR: ultrafast universal RNA-seq aligner. Bioinformatics. Jan 1 2013;29(1):15–21. doi:10.1093/bioinformatics/bts635

98. Love MI, Huber W, Anders S. Moderated estimation of fold change and dispersion for RNA-seq data with DESeq2. Genome Biol. 2014;15(12):550. doi:10.1186/s13059-014-0550-8

99. Huber W, et al. Orchestrating high-throughput genomic analysis with Bioconductor. Nat Methods. Feb 2015;12(2):115–21. doi:10.1038/nmeth.3252

100. Mootha VK, et al. PGC-1alpha-responsive genes involved in oxidative phosphorylation are coordinately downregulated in human diabetes. Nat Genet. Jul 2003;34(3):267–73. doi:10.1038/ng1180

101. Langmead B, et al. Ultrafast and memory-efficient alignment of short DNA sequences to the human genome. Genome Biol. 2009;10(3):R25. doi:10.1186/gb-2009-10-3-r25

102. Zhang Y, et al. Model-based analysis of ChIP-Seq (MACS). Genome Biol. 2008;9(9):R137. doi:10.1186/gb-2008-9-9-r137

103. Schep AN, et al. chromVAR: inferring transcription-factor-associated accessibility from single-cell epigenomic data. Nat Methods. Oct 2017;14(10):975–978. doi:10.1038/nmeth.4401

104. motifmatchr: Fast Motif Matching in R. R package version 1.12.0; 2020.

105. Liao Y, Wang J, Jaehnig EJ, Shi Z, Zhang B. WebGestalt 2019: gene set analysis toolkit with revamped UIs and APIs. Nucleic Acids Res. Jul 2 2019;47(W1):W199-W205. doi:10.1093/nar/gkz401

